# Unravelling drug-resistant proteotypes through phenotype-resolved proteomics of single-cell derived colonies

**DOI:** 10.1101/2025.10.11.681009

**Authors:** Di Qin, Ridhima Das, Simon Schallenberg, Kristoffer Riecken, Ingeborg Tinhofer, Fabian Coscia

## Abstract

Drug resistance remains a major cause of treatment failure and disease progression and is closely linked to intratumoral heterogeneity. Mass spectrometry (MS)-based single-cell proteomics (SCP) can resolve drug-resistant cell states, but current methods lack clonal and phenotype information and are often confounded by cell-cycle and cell-size effects. Here, we introduce PhenoSCoP, a microscopy-guided discovery proteomics framework that profiles single-cell-derived colonies to map proteomic heterogeneity while preserving clonal origin and image-defined phenotypes. In head and neck squamous cell carcinoma models, PhenoSCoP distinguished heritable clone-intrinsic cisplatin (CP) response programs from adaptive treatment-induced changes, uncovered pre-existing resistant-like proteotypes in treatment-naïve populations, and identified an NDUFB11-associated mitochondrial vulnerability in a resistant clone. In patient-derived primary HNSCC cells, CP-tolerant cells resolved into two proteomic states with distinct phenotypes and high or low S100A7/A8/A9 expression. These programs were also detected in HNSCC patient tissues and, in an illustrative matched progression case, increased across metastatic sites. PhenoSCoP therefore provides a phenotype-resolved framework for discovering stable and adaptive proteotypes associated with therapeutic response.

## Introduction

Resistance to anti-cancer therapies, including targeted therapies, chemotherapies, and immunotherapies, remains a major obstacle for successful treatment outcomes ^1^. While extensive research has focused on the genetic mechanisms of drug resistance and tumor heterogeneity ^2,3^, non-genetic mechanisms that confer transient or persistent resistance are increasingly being recognized as important drivers of therapeutic responses ^4^. Cancer cells exploit both genetically-encoded and adaptive drug resistance programs, emphasizing the importance of methodologies that capture both mechanisms on a global, quantitative, and phenotype-resolved scale ^5^. Studying the global proteome of distinct tumor cell clones, a close proxy for cellular function ^6^, is therefore of paramount importance to unravel the mechanisms underlying differential treatment responses and identify clone-specific vulnerabilities.

A growing body of literature supports the notion that cellular states before drug treatment dictate treatment outcomes ^7–9^, including observations from clinical samples ^10,11^. Dissecting such pre-existing molecular programs associated with clone-specific survival is hence of high clinical value for discovering novel therapeutic strategies and for eradicating the most aggressive and resistant clones within heterogeneous tumors. However, the extent to which clonal proteomic heterogeneity and pre-existing proteotypes drive therapeutic outcomes and drug resistance remains poorly understood. Based on imaging-based proteomics, it has been estimated that one-fifth of the human proteome exhibits cell-to-cell variability ^12^, emphasizing the importance of single-cell analyses in the context of treatment responses. High-content imaging offers detailed insights into the proteome at subcellular and spatiotemporal resolutions, yet only a few proteins can be profiled simultaneously ^13–16^. Mass spectrometry-based single-cell proteomics (SCP) offers a complementary strategy and currently achieves a depth of a few thousand proteins ^17–21^. Nevertheless, SCP remains challenging to interpret because of incomplete proteome coverage, missing values, and strong confounding effects arising from differences in cell size and cell-cycle state ^22–24^. These limitations motivate hybrid approaches that combine imaging with deep discovery proteomics while retaining information about clonal origin.

To address this gap, we propose PhenoSCoP, a microscopy-guided discovery proteomics concept that combines fluorescence imaging, laser microdissection (LMD), and ultrasensitive MS-based proteomics. Rather than profiling individual cells, we developed a scalable workflow to profile single-cell-derived colonies without extensive serial dilution and clonal expansion experiments. Inspired by elegant studies on transcriptional memory, persisting over several cell cycles, ^25^ weeks, ^8^ or even years ^26^, we hypothesized that single-cell-derived colonies retain distinct and quantifiable proteomic traits that contribute to stable drug-response phenotypes. Although protein-level memory and its functional consequences have been demonstrated previously ^27,28^, the extent to which the cellular proteome features heritable, clone-specific, and phenotype-defining traits remains largely unexplored. Our colony-centric approach prioritizes heritable protein-level changes while mitigating short-lived and transient fluctuations that often confound SCP experiments.

Using head and neck squamous cell carcinoma models, we show that PhenoSCoP can distinguish stable clone-intrinsic proteomic programs from treatment-induced adaptive responses. Our approach also uncovered rare pre-existing proteotypes within drug-naïve populations, identified clone-specific differences in proliferation and cell-cycle dynamics, and enabled matched proteomic analysis of the same colonies before and after CP exposure through a live-cell colony-splitting strategy. PhenoSCoP was also successfully applied to patient-derived primary cells, where it identified several of the same proteins and pathways associated with CP tolerance in the cell line model. Analysis of patient tumor proteomes further showed that these resistance-associated proteins were detectable and variable across patients and spatially distinct disease sites. Together, PhenoSCoP provides a scalable strategy for linking clonal phenotypes to stable and adaptive proteomic programs associated with therapeutic response.

## Results

### An optimized workflow for immunofluorescence microscopy guided discovery proteomics

Our goal was to develop an imaging-guided discovery proteomics method that captures clone-specific proteotypes that are functionally linked to drug responses (**Figure 1A**). We devised **Pheno**type-resolved **S**ingle-**Co**lony **P**roteomics (PhenoSCoP), which combines clonogenic cell growth, fluorescence microscopy, LMD and ultrasensitive mass spectrometry (MS)-based proteomics. Our method was inspired by the clonogenic survival assay first described in 1956 by Puck and Marcus ^29^. This assay is a gold standard for evaluating cellular viability and measuring the proportion of surviving reproductive cells following treatment, thereby enabling the investigation of long-term cellular outcomes ^30^. Instead of analyzing single cells, we sought to analyze single-cell-derived phenotypes. Cells are seeded on microscopy slides at low concentrations, exposed to drug treatment, and outgrowth into multicellular colonies is monitored, visualized and quantified. Finally, drug-tolerant and colony-forming cells are isolated and their proteomes compared to untreated controls.

**Figure 1.**
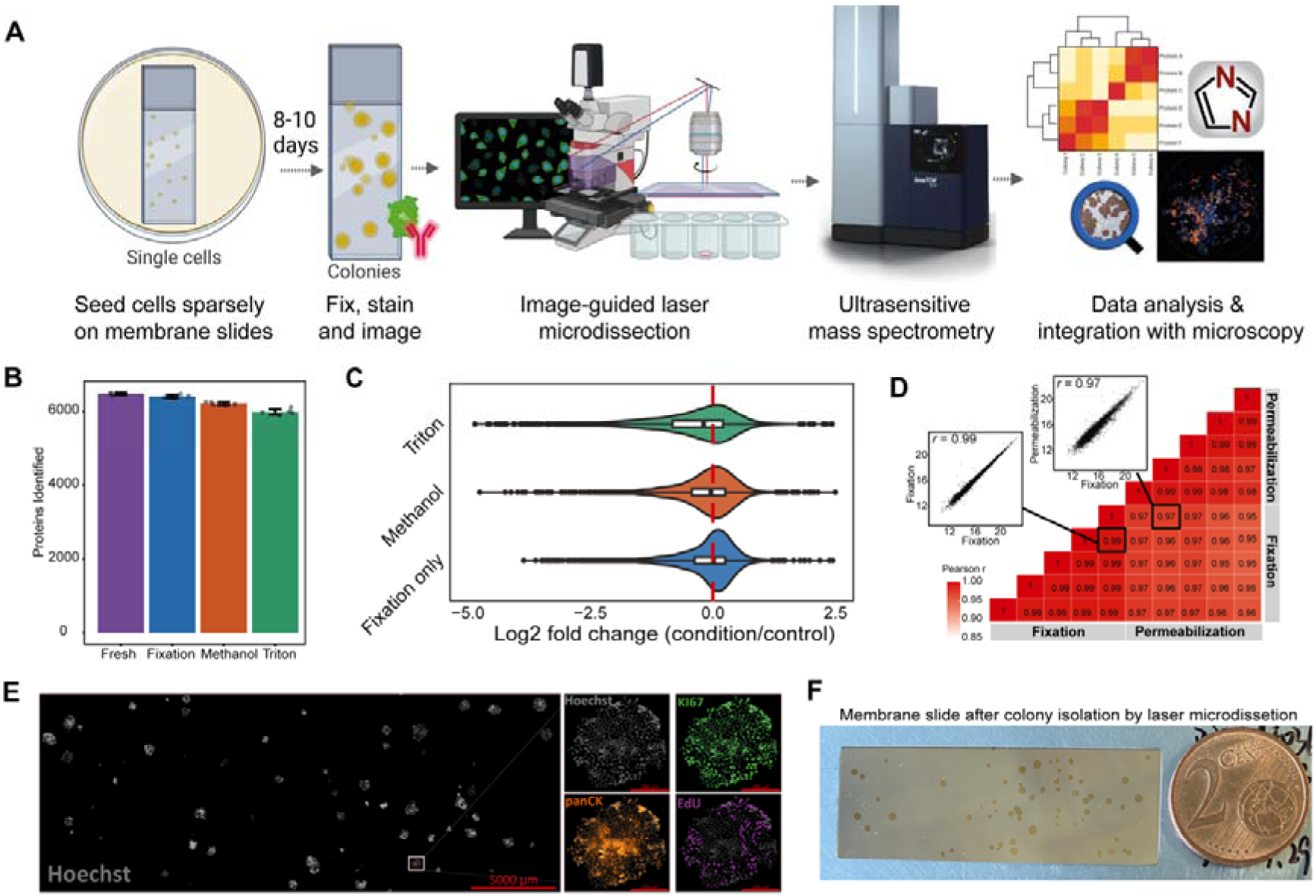
Optimization and proteomic validation of the PhenoSCoP workflow. (A) Schematic representation of the PhenoSCoP approach. (B) Comparison of protein identifications from approximately 100 laser microdissected cells prepared using four different protocols: fresh cells, formaldehyde (FA) fixed cells, FA fixed cells permeabilized using ice-cold methanol or Triton-X-100. (C) Quantitative proteomic comparison of fixation-only and permeabilized cells relative to untreated (fresh) control cells. Only common proteins were used without data imputation. (D) Sample correlation matrix (Pearson’s *r*) of al l replicates for fixation-only and methanol-permeabilized cells. (E) Whole-slide immunofluorescence imaging of colonies grown on a PPS membrane slide using the optimized staining protocol. One representative colony corresponding to the white box is shown in the right panel, highlighting nuclear and cytosolic staining. Hoechst (grey), Ki-67 (green), Pan-CK (orange) and EdU-positive S-phase cells (pink). Scale bar for the overview image (left panel) is 5,000 µm, and 200 µm for example colonies (right panel). (F) Brightfield image of the PPS slide after colony isolation using laser microdissection.

To realize PhenoSCoP, we first optimized the staining and imaging workflow and assessed the impact of fixation and permeabilization on the quantitative proteomic read-out. FaDu cells, an established polyclonal model for studying CP resistance in HNSCC ^31^, were seeded on polyphenylene sulfide (PPS) metal frame slides required for laser microdissection, grown to ∼80% confluence, and subjected to different protocols prior to LMD and MS-based proteomics (**Figure 1B**). The comparison of fresh FaDu cells with formaldehyde-fixed cells (4%, 10 min) resulted in almost identical proteome coverage, in line with a recent study ^32^. From the small 100-cell regions, we quantified more than 6,000 unique proteins in a 15-min active liquid chromatography (LC) gradient on a Bruker timsTOF Ultra instrument operated in diaPASEF mode ^33^ and based on library-free DIA-NN ^34^ analysis (**Figure 1C**). To enable the staining of intracellular proteins critical for image-based cell phenotyping, we tested two standard permeabilization protocols based on 90% ice-cold methanol or 0.2% Triton X-100. While both protocols resulted in slightly lower proteome coverage than fixation alone (< 4%), quantitatively, the methanol protocol performed better and yielded results more comparable to the fixation-only condition than Triton X-100 treated cells (**Figure 1C, Table S1**), in good agreement with previous work ^35^. Importantly, despite the lower number of identified proteins after permeabilization, global proteomic inter-group correlations were still excellent (median Pearson *r* = 0.97, **Figure 1D**) and yielded similar cellular compartment distributions (**Figure S1A**). Therefore, formaldehyde fixation combined with methanol permeabilization was selected for our final sample preparation protocol. An additional comparison of cells grown on PPS membrane slides versus standard culture plates revealed comparable colony growth and highly similar quantitative proteomic profiles (**Figure S1B-C**). Next, we seeded FaDu cells onto PPS metal frame slides and monitored colony formation over ten days of growth. At day ten, cells were formaldehyde-fixed and subjected to whole-slide immunofluorescence imaging (**Figure 1E**). We validated the high staining and imaging quality using primary conjugated antibodies targeting cytoplasmic pan-cytokeratin (panCK) and the proliferation marker Ki-67, in addition to EdU (marker of active DNA synthesis in S-phase) and DAPI (DNA) (**Figure 1E**). Following imaging, we isolated distinct colonies by laser microdissection, providing sufficient cellular input for deep and robust proteome profiling (**Figure 1F**).

### PhenoSCoP effectively captures clonal differences while mitigating SCP confounders

We aimed to develop a method that captures single-cell-derived phenotypes while minimizing SCP confounders, particularly cell-cycle-associated variation in protein abundance, cell size, and proteome coverage ^36^ (**Figure 2A**). We reasoned that profiling multicellular colonies would reduce the influence of these transient single-cell effects, while imaging preserves colony-level cell-cycle information and the proteomic readout prioritizes more persistent, clone-associated differences.

**Figure 2.**
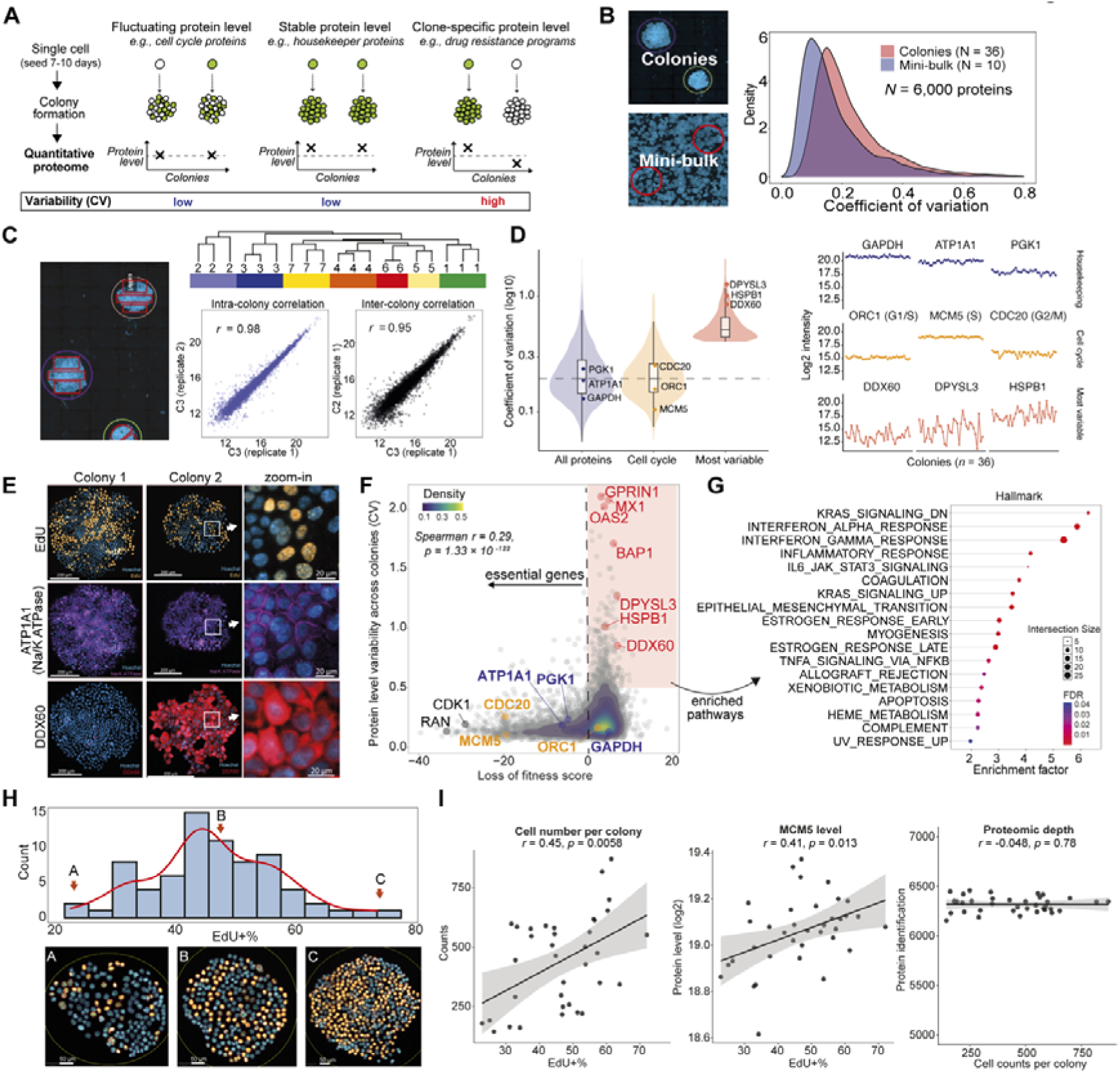
PhenoSCoP captures clone-specific proteomic signatures while mitigating SCP confounders. (A) Schematic scenarios of protein level variation in single-cell-derived colonies. Fluctuating, stable, and hereditary protein-level changes are distinguished from quantitative proteome measurements. (B) Left: Example images of Hoechst-stained colonies and homogenously grown cells (mini-bulk). Right: Density plot showing the coefficient of variation (CV) of protein quantification from single-cell-derived colonies (n = 36) and mini-bulk replicates (n = 10). Mini-bulk samples were collected from homogeneously seeded cell regions (∼400,000 µm²). Colony injections were normalised to 178 ± 11 cells (mean ± SD) based on image-derived cell counts. (C) Intra-colony proteomic comparison. Seven colonies were separated into two to three segments using laser microdissection and subjected to proteome analysis. Unsupervised hierarchical clustering based on 6,088 quantified proteins. Sample correlations between segments from the same colony (intra-colony) and different colonies (inter-colony) are shown. (D) (Left) Distributions of colony-to-colony coefficients of variation (CV) for quantified proteins. CV was calculated on linear intensities across 36 colony samples for proteins detected in ≥50% of colonies. Violin plots show (i) all proteins (n = 6,168), (ii) Human Protein Atlas cell-cycle-dependent (CCD) proteins ^12^ detected in the dataset (n = 320), and (iii) the most variable proteins (top 10% CV; n = 617). Inner boxplots indicate the median and interquartile range; the dashed line marks the proteome-wide median CV (0.195). Labeled points highlight housekeeping proteins (PGK1, ATP1A1, GAPDH) on the all-protein distribution, selected cell-cycle markers (ORC1, MCM5, CDC20) on the CCD distribution, and highly variable proteins (DDX60, DPYSL3, HSPB1) on the most-variable distribution. CV is shown on a log₁₀ scale. (Right) log₂ intensity profiles of the same nine proteins across the 36 colonies (shared y-axis). (E) Immunofluorescent images of representative markers from different colonies, corresponding to the three protein types shown in panel d. Scale bar is 200 µm for colonies and 20 µm for the indicated regions. (F) Protein variability (coefficient of variation) versus gene essentiality (loss of fitness score in FaDu cells). Proteins from panel d are highlighted: yellow (cell cycle), blue (housekeeping), and red (variable proteins). Cell-cycle and housekeeping proteins generally showed lower variability and stronger dependency. (G) Hallmark pathway enrichment of non-essential variable proteins from panel f. (H) Upper panel: Histogram showing the proportion of S-phase cells across 73 colonies determined by counting EdU-positive cells from I F image data. Lower panel: Representative images of colonies with low, medium, and high proportions of S-phase cells, corresponding to the upper panel. Image analysis was implemented in QuPath. (I) Scatter plots showing the correlations of cell numbers (left) and MCM5 protein abundance (log₂ intensity, middle) with the proportion of EdU-positive (EdU+) cells, and cell counts per colony with the number of protein identifications (right) across individual colonies derived from FaDu parental cells. Linear regression lines with 95% confidence intervals are shown.

To test whether our approach effectively captured clone-related proteotypes (**Figure 2A**), we applied PhenoSCoP to the polyclonal HNSCC model FaDu. We isolated and profiled 36 colonies and 10 mini-bulk samples acting as controls. Mini-bulk samples consisted of homogenously seeded cells that did not grow into individual colonies but were otherwise cultivated under identical growth conditions (**Figure 2B**). All samples were processed and measured within the same experimental batch. On average, we consistently quantified 6,208 proteins per colony with high data completeness across samples, which was, as expected, much superior to single-cell proteomic measurements of the same cells (**Figure S2A-B**). This allowed us to successfully capture biological differences between individual colonies, as proteomic variation from different colonies significantly exceeded that of the mini-bulk measurements (median CV 20% *vs*. 14%, p < 2.2e-16) (**Figure 2B**). In support of this, the proteomes of cells isolated from the same colony were quantitatively more similar and grouped together in unsupervised hierarchical clustering compared to cells from neighbouring colonies (**Figure 2C**). Together, these results indicated that our approach captured hereditary proteomic traits shared between cells of the same subclone.

Next, we reasoned that our precise protein quantification strategy in combination with colony average measurements would also allow us to distinguish different biological sources of protein-level variability. Proteins with canonical housekeeping functions (e.g., GAPDH,

ATP1A1, and PGK1) showed low colony-to-colony variability, with CVs near or below the proteome-wide median (**Figure 2D**). In contrast, a large subset of proteins displayed marked abundance differences across single-cell-derived colonies. Among the most variable were the metastasis-associated protein DPYSL3 ^37^, the interferon-response factor DDX60, and heat shock protein beta-1 (HSPB1). Cell-cycle related proteins such as ORC1, MCM5, and CDC20 showed comparably low variation. Overall, we found an inverse correlation between protein level variability and protein essentiality, as revealed by the integration of the cancer dependency map (DepMap) dataset ^38^ (**Figure 2F, Table S2**). In other words, the most variable proteins tended to be less essential (Spearman r = 0.29, p = 1.33 × 10⁻¹²²), whereas cell-cycle-dependent (CCD) proteins ^12^ and ‘housekeeping’ proteins on average showed lower variation, but higher dependency scores (**Figure 2F and Figure S2C**). Immunofluorescence microscopy confirmed the different protein level patterns revealed by PhenoSCoP. For example, while DDX60 showed variable and colony-specific expression (**Figure 2E**), ATP1A1 (Na/K ATPase) staining was positive for the majority of cells independent of the colony. We found that the most variable and non-essential proteins were strongly enriched for cancer and immune-related pathways, such as interferon response, KRAS signalling, and epithelial-to-mesenchymal transition (**Figure 2G**).

The seamless integration of imaging and proteomic data further allowed us to assess proliferation and cell cycle differences between colonies by comparing colony size (imaging), the fraction of cells in S phase (EdU labeling), and cell cycle protein abundance measurements (proteomics) (**Figure 2H-I**). Nearly half of all cells (47% ± 11%) were in early or late S-phase across colonies. However, we also identified clear outlier phenotypes that deviated from this cell line average. For example, one colony mostly featured slow-cycling cells characterized by a lower total cell number and only 25% of EdU+ cells, whereas two colonies were more proliferative resulting in higher cell count and 70% EdU+ cells (**Figure 2H**). While colony size and replication-related protein levels (e.g., MCM5) were generally positively correlated with the EdU+ cell fraction (Spearman *r* = 0.45 and 0.41, respectively), several more proliferative colonies showed a low fraction of EdU+ cells (**Figure 2I**), indicative of clonal differences. Importantly, irrespective of proliferation status, cell cycle state or colony size, proteome coverage was highly comparable across the analyzed phenotypes (Figure 2I). Hence, our approach provides a robust and balanced solution to profile clone-specific and rare cellular phenotypes while integrating cell cycle information.

### PhenoSCoP identifies molecular drivers of cisplatin resistance

Cisplatin (CP) is a cornerstone chemotherapy drug used to treat HNSCC, particularly in advanced stages; however, treatment failure due to primary and acquired resistance is common, particularly in human papilloma virus (HPV)-negative carcinomas ^39^. High intratumoral heterogeneity and expansion of pre-resistant clones are the main drivers of CP resistance in HNSCC ^40–42^. The HPV-negative FaDu cell line is an established model for studying CP resistance in HNSCC and consists of multiple genetically and transcriptionally distinct subclones with varying CP sensitivities, differing by more than 100-fold ^43^. In this cell line, Niehr *et al*. revealed a causal relationship between the *TP53* p.R248L gain-of-function (GOF) missense mutation and increased CP resistance. Importantly, stabilizing *TP53* GOF mutations are also associated with poor patient outcomes ^44–46^, emphasizing the need to better understand the molecular underpinnings and therapeutic vulnerabilities of such aggressive clones. To test whether our method could identify pre-existing and adaptive proteotypes associated with CP resistance, we employed a multilayered approach. First, we applied PhenoSCoP to characterize phenotypic and proteomic differences between two previously established single-cell-derived FaDu subclones of known CP sensitivity (Figure 3A-E). Second, we derived a live-cell colony-splitting assay to obtain matched proteomic data from before and after drug exposure of the same subclone to investigate adaptive treatment responses (Figure 3F-J). Lastly, we asked whether chemotherapy resistance-associated proteotypes could also be identified directly from treatment-naïve parental cells using our approach (Figure 4).

**Figure 3.**
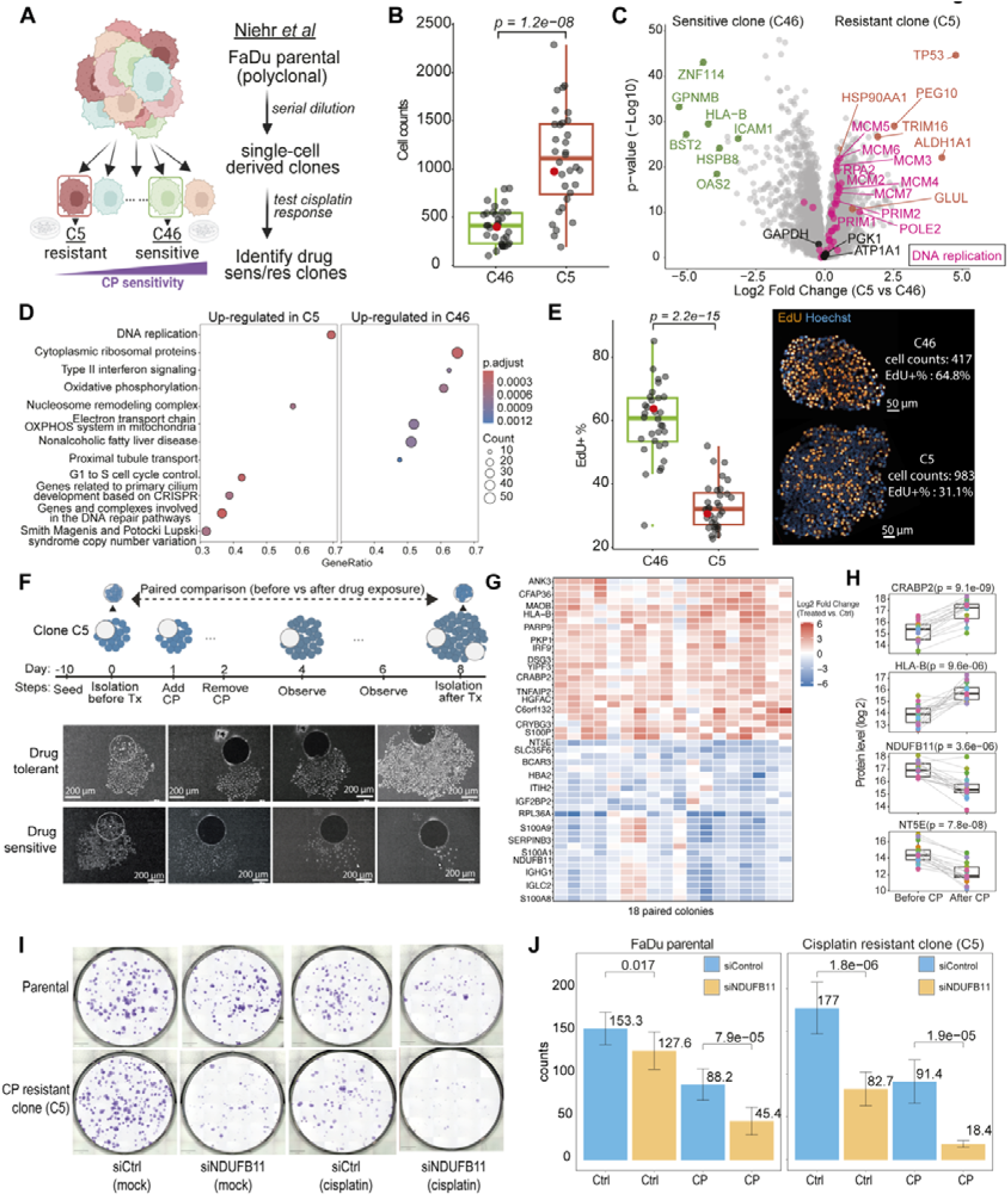
PhenoSCoP resolves stable and adaptive proteomic programs associated with CP resistance. (A) Schematic representation of two previously characterized drug-naïve clones isolated from the FaDu parental cell line. Clone C5 showed higher primary CP resistance than clone C46. (B) Boxplot of cell counts per colony for C46 (n = 32) and C5 (n = 32). Colony sample injection amounts were normalized to an average of 237 ± 47 cells (mean ± SD) per measurement using image-derived cell counts. Red dots indicate the colonies shown in the representative images in the right panel of Fig. 3E. (C) Volcano plot comparing protein expression between C5 and C46 single cell-derived colonies. Stable housekeeping proteins (GAPDH, PGK1, and ATP1A1) are shown in black. Proteins belonging to the DNA replication pathway (WikiPathways) are highlighted in pink. Selected top differentially expressed proteins are labeled, with proteins upregulated in C5 shown in orange and those upregulated in C46 shown in green. (D) Enriched pathways (Wiki pathways) comparing C5 and C46 colonies to identify biological differences between clones. (E) Left: Boxplot of the proportion of S-phase cells in C46 and C5 colonies, quantified as the percentage of EdU-positive (EdU+) cells from immunofluorescence (IF) images. Red dots indicate the colonies shown in the representative images in the right panel. Right: Representative IF images of C46 and C5 colonies showing EdU staining. The total cel number and percentage of EdU+ cells are indicated alongside each image. (F) Top: Schematic diagram and representative images of the colony-splitting experiment using the C5 clone. 80,000 - 100,000 µm² were isolated from colonies before and after drug exposure. Bottom: Representative Images of colonies before and after LMD. Scale bar = 200 µm. (G) Heatmap of the top 50 shared upregulated and downregulated genes identified across 18 paired colonies before and after CP exposure. (H) Boxplots showing representative proteins identified by paired colony analysis as being consistently altered following CP treatment (n = 18 paired colonies). Each dot represents an individual colony, and lines connect matched before- and after-treatment samples. Colours indicate colony id. (I) Representative images from clonogenic assays showing the effects of NDUFB11 knockdown and CP treatment on both parental and resistant (C5) FaDu cells. (J) Quantification of colonies from knockdown experiments shown in panel I. Data represent the average number of colonies from three independent experiments. Significance was assessed by a two-sided Student’s t-test.

**Figure 4.**
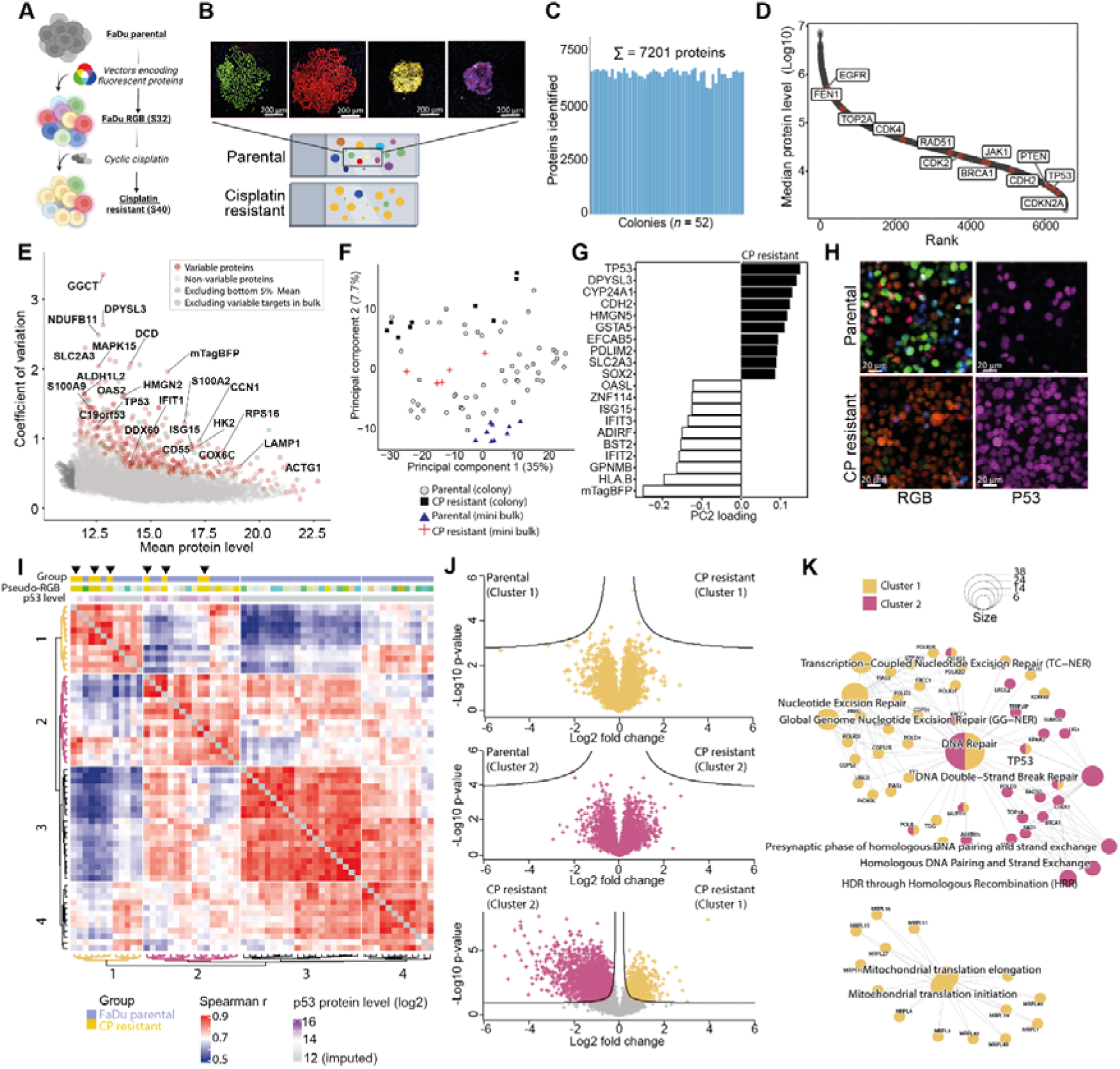
PhenoSCoP identifies pre-existing proteotypes driving primary CP resistance. (A) Schematic of the lentiviral vector-mediated RGB-marked FaDu model system. (B) Representative images of colonies derived from FaDU RGB clones. Scale bar: 200 µm. (C) Depth of 52 colony proteomic measurements. (D) Dynamic range of protein abundance from colony proteome measurements, with examples of oncogenes and tumor suppressor proteins highlighted. (E) Comparison of protein level variability (CV) and mean protein levels of all identified proteins. Variable proteins (red) are defined as the top 5% of the most variable proteins per binned segment. One hundred bins were used to cover the entire protein level range. Proteins with high variation in pseudo-bulk samples were excluded. Mini-bulk samples were collected from homogeneously seeded cell regions (∼305,000 µm²). Colony injections were normalised to 276,000 ± 60,000 µm² (mean ± SD) based on colony area. (F) Principal component analysis (PCA) of FaDu primary clones (n = 52), primary mini-bulk samples (n = 9), resistant clones (n = 9), and resistant mini-bulk samples (n = 5) using 357 variable proteins. (G) Top 20 features contributing to principal component two of PCA in panel F. (H) Immunofluorescence staining of TP53 in primary and resistant FaDu cells. Scale bar: 20 µm. (I) Proteome correlation analysis based on the most variable proteins (related to panel E, filtered for 70% valid values). Distances were calculated using Euclidean metrics and hierarchical clustering using average linkage, revealing four main clusters. (J) Volcano plots comparing CP-resistant versus drug-naive colonies from Cluster 1 (top) and Cluster 2 (middle). Bottom: Volcano plot comparing CP-resistant colonies from Cluster 1 versus Cluster 2. Cluster information was extracted from panel I. (K) Enriched Gene Ontology pathways of proteins significantly upregulated in CP-resistant clones (panel I and J, Clusters 1 and 2) compared to Clusters 3 and 4, which included drug-naïve proteomes of FaDu parental colonies.

We first profiled the proteomes of two single-cell-derived and drug-naive clones representing stable CP-sensitive (C46 clone) and resistant (C5 clone) phenotypes (Figure 3A, Table S3). Although both clones had an intronic *TP53* mutation (*TP53* c.673 G>A) resulting in an early stop codon, the CP-resistant C5 clone carried an additional *TP53^R248L^* GOF mutation and an extra copy on chromosome 17 (encoding mutant p53). Additional mutations are summarized in Figure S3A. To confirm the stability of different levels of CP resistance, we combined PhenoSCoP with CP treatment. Cells from both clones were plated as single-cell dilutions on PPS membrane slides and treated for 24 hours with their expected IC-50 CP concentrations (C46:30 ng/mL, C5:150 ng/mL). On day 10, the colonies were fixed, Hoechst stained, and subjected to whole-slide imaging and global proteomic analysis. CP treatment resulted in a reduction of approximately 50% in colony formation for both clones, confirming their respective IC-50 dosages (Figure S3B). The resistant clone showed substantially greater net expansion, forming colonies up to threefold larger over the same growth interval (Figure 3B). Proteomic analysis revealed extensive quantitative differences between the two subclones, with 968 differentially abundant proteins. Although individual DNA-replication and proliferation-associated proteins showed relatively modest fold changes, their highly consistent and coordinated regulation resulted in a clear enrichment of DNA-replication processes in the resistant clone (FDR < 0.05; Figure 3C-D). Consistent with our previous findings (Figure 2D), non-significant proteins included ‘housekeeping’ proteins, such as GAPDH, ATP1A1, and PGK1. Notably, p53 was the most upregulated protein in the CP-resistant clone (Figure 3C), highlighting the strong protein stabilization effect of the R248L GOF mutation and its functional role as a driver of CP resistance ^42^. The previously reported amplification of chromosome 17, encoding *TP53*, was also strongly reflected in the quantitative proteome of this clone (Figure S3D). In the sensitive clone, p53 was undetectable at baseline (Figure S3E), in agreement with the destabilizing intronic *TP53* mutation. Our exploratory proteomic data also shed light on the potential cellular mechanism underlying p53 stabilization. We identified the chaperone HSP90 (HSP90AA1), a critical mediator of mutant p53 stabilization and localization ^47,48^, among the top significantly regulated proteins (Figure 3C). Pathway enrichment analysis revealed upregulated DNA damage repair, ribosomal processes, and p53 signaling, whereas the sensitive clone showed higher levels of proteins related to antigen presentation, interferon, and TNF signalling (Figure 3D). The concurrent enrichment of DNA damage repair and proliferation-associated signatures in the resistant clone was particularly intriguing. While elevated DNA repair capacity could provide a plausible mechanism for the five-fold higher CP tolerance, the pronounced proliferative phenotype (Figure 3B-C) initially appeared counterintuitive, since platinum agents preferentially affect proliferating cells. However, our immunofluorescence imaging-grounded approach helped to resolve this by discovering that the resistant clone had significantly reduced S-phase occupancy (Figure 3E). Together, these data indicate that stable resistance of the *TP53^R248L^* GOF-driven clone combines enhanced DNA damage repair capacity and altered cell-cycle dynamics, in which faster colony expansion coexists with a lower fraction of cells undergoing DNA synthesis at any given time.

Together, these findings defined stable, clone-intrinsic proteomic programs that are consistent with the genetically encoded differences between the resistant and sensitive clones. However, as resistance not only arises from genetically driven mechanisms, but also from adaptive and non-genetic programs, we next investigated whether PhenoSCoP could also reveal CP-induced adaptive responses. To this end, we developed a live-cell colony-splitting approach to obtain matched proteomic data from the same colonies before and after drug exposure. This strategy enables direct measurement of adaptive proteome changes within identical colony backgrounds from CP-resistant (C5) phenotypes. From 21 colonies that we monitored over eight days in culture, we identified 18 as drug tolerant that continued expanding subsequently after CP removal (Figure 3F).

Proteins altered in drug-tolerant colonies were enriched in processes previously associated with non-genetic resistance (Figure S3F-G). For example, constitutive type-I-interferon signaling has been identified as a non-genetic driver of primary drug resistance in several cancers ^7,9,49,50^. We also identified several proteins involved in mitochondrial metabolism and OXPHOS that showed differential abundance (Figure S3F-G). NDUFB11, a complex I mitochondrial protein, was of particular interest, as it was among the most downregulated proteins after CP treatment (Figure 3G-H) and has recently been associated with enhanced CP resistance in ovarian cancer models through increased protein degradation ^51^. This led us to hypothesize that dynamic regulation of NDUFB11 contributes to CP response. To test whether NDUFB11 was indeed functionally relevant for CP tolerance in the FaDu model, we performed siRNA-mediated NDUFB11 knockdown using 3 nM siRNA with and without CP. This resulted in a marked decrease in colony formation in the resistant C5 clone (−53%), whereas the polyclonal parental line was much less affected (−17%) (Figure 3I-J). Notably, when combined with CP treatment, colony formation was nearly abolished in the resistant clone. Although NDUFB11 is classified as non-essential in most cancer cell lines, CRISPR-based knockout data indicated a moderate dependency in FaDu cells (Figure S3H). Our results suggest that this dependency could be clone-specific and might be substantially greater in the p53-mutant C5 clone. Further research is needed to determine whether targeting NDUFB11, or more globally mitochondrial redox metabolism, could be exploited therapeutically to eradicate p53-mutant HNSCC clones.

Collectively, these data demonstrate the potential and versatility of PhenoSCoP to distinguish stable clone-intrinsic proteomic programs from dynamic, non-genetic adaptations and to prioritize functionally relevant drivers of therapeutic responses.

### PhenoSCoP identifies pre-existing proteotypes driving primary CP resistance

Increasing evidence suggests that the cellular state prior to drug administration plays a crucial role in determining therapeutic responses ^7,8,11^. Therefore, we next investigated whether PhenoSCoP could reveal proteotypes associated with primary drug resistance. As both the C46 and C5 clones were derived from treatment-naïve FaDu parental cells, these proteotypes served as excellent ground truth to be picked up by our method. To support clonal tracking and exclude colonies arising from overlap or fusion events, we integrated fluorescence-based RGB labeling ^52,53^. Following whole-slide imaging, we isolated 52 spatially distinct colonies displaying non-mixed RGB patterns. Through stochastic color mixing after lentiviral expression of different fluorescent proteins in varying but stable amounts, the stably inherited RGB profile provided a visual clonal-tracking label, although similar color profiles may occur independently. This approach enabled us to monitor clonal selection after long-term cyclic CP treatment and assess whether drug-resistant proteotypes were already pre-existing or developed through proteome remodelling. To this end, FaDu parental cells were stably transduced with RGB vectors, enabling the visualization of diverse clonal populations (Figure 4A). Following whole-slide RGB imaging (Figure 4B), we isolated 52 color-unique colonies and quantified over 7,200 proteins in total, with a median of more than 5,800 proteins per colony (Figure 4C). Our dataset included many known oncogenes and tumor suppressors involved in HNSCC carcinogenesis, such as EGFR, TP53, BRCA1, and CDH2 (Figure 4D). To systematically identify the most variable fraction of the proteome across single-cell-derived colonies, we calculated the coefficient of variation (CV) for each protein and plotted them against the mean protein abundance (Figure 4E, Table S4). To avoid potential bias from low-abundance proteins, which generally exhibit higher variability, we binned the data into 100 segments and retained only the top 5% of each segment. Additionally, the 5% lowest-abundant proteins and those showing high variation in our mini-bulk controls were excluded (Figure 2B,D). Using this strategy, we prioritized 357 variable proteins with high confidence. The protein sequence coverage of the selected proteins was similar to that of the full dataset (Figure S4A), further supporting the idea that our approach prioritized true biological and clone-specific signals rather than technical noise. An independent replicate dataset also confirmed a substantial overlap among the prioritized proteins, which was more pronounced at the pathway level (Figure S4B, Table S4). Using this approach, we not only captured C5 and C46 clone-specific signatures, such as p53, DPYSL3, and the interferon-response proteins HLA-B, IFIT1, and DDX60 (Figure 4E), but also the quantified RGB reporter peptides (e.g., mTagBFP), which showed the expected variability and confirmed that different clones were compared. Cell cycle-dependent proteins ^12^ instead exhibited only low colony variation, consistent with their more transient fluctuation (Figure S4C) and our previous results (Figure S2C). Repeated doses of CP led to strong clonal selection, as was evident from a dominant orange clone revealed by fluorescence imaging (Figure S4D). Interestingly, principal component analysis closely grouped drug-resistant clones with a small fraction of colonies derived from the treatment-naïve parental population. This finding was obscured when we compared the proteomes of clonal mixtures (mini-bulk controls) (Figure 4F), underlining our method’s resolution in identifying clone-specific molecular programs. P53, DPYSL3, CYP24A1, and N-cadherin (CDH2) were the most strongly elevated in these pre-resistant cells (Figure 4G-H). Proteome correlation analysis further showed that the resistant proteotypes were highly related to a few drug-naïve parental clones (Pearson *r* > 0.9, Figure 4I). While immunofluorescence (IF) imaging and RGB pseudo-color inference from reporter peptide quantification (Figure S4E) suggested that one dominant yellow/orange clone survived multiple rounds of CP (Figure 4H-I), our proteomic data suggested two distinct proteotypes of similar color profile (Figure 4I-K). Although this observation could possibly be explained by one pre-resistant clone that underwent additional proteome remodelling after cyclic CP exposure, comparison with untreated cells of the same color-matched subcluster revealed remarkably similar proteomes with no significant differences (Figure 4J). Both resistant clones in clusters 1 and 2 showed high mutant p53 levels and DNA damage repair signatures (Figure 4I and 4K), but with nuanced differences. While the resistant clone of cluster 1 was characterized by higher protein levels related to mitochondria and nucleotide excision repair, the main pathway for detecting and repairing CP-induced DNA adducts ^54,55^, the cluster 2 clone showed higher homologous recombination-based DNA damage repair (Figure 4K), suggesting different modes of DNA damage repair.

Together, these data show that PhenoSCoP can resolve rare, pre-existing proteomic states within treatment-naïve populations that closely resemble subsequently selected cisplatin-resistant clones. This highlights the ability of colony-resolved proteomics to distinguish pre-existing resistance-associated proteotypes from treatment-induced remodeling that would be obscured in population-averaged measurements.

### PhenoSCoP resolves heterogeneous CP-tolerant states in primary HNSCC cells

Having established that PhenoSCoP can uncover clone-specific CP-resistant proteotypes in an established HNSCC cell line, we next asked whether the same phenotype-resolved workflow could be applied to more clinically relevant models. We therefore applied PhenoSCoP to patient-derived primary HNSCC cells (Figure 5A). Primary cells were seeded sparsely on membrane slides, treated with CP for 24 h and, after drug washout, clonogenic survival was monitored over ten days to isolate drug-tolerant colonies alongside untreated controls. EdU labelling showed that cells that survived the treatment recovered their proliferative capacity (Figure S5A). As expected, untreated colonies were much larger, confirming the strong anti-proliferative effect of CP (Figure 5B). To account for colony-size differences before proteomics, we normalized sample injection to 50-cell equivalents from the imaging-derived total colony area, yielding comparable proteome coverage across groups and phenotypes (Table S5, median 5,537 protein groups per colony) with high quantitative reproducibility (median Pearson *r* = 0.97, Figure S5B-C). Principal component analysis of all colony proteomes separated the two conditions along PC1 (Figure 5C). CP-tolerant colonies were strongly enriched for the calcium-binding proteins S100A7/A8/A9, linked to epithelial differentiation and HNSCC progression via NF-κB and PI 3K/Akt/mTOR signalling ^56,57^, together with cornification- and squamous differentiation-associated proteins such as CALML5, SERPINB3, DSC1 and CASP14 (Figure 5D). Interestingly, CP-tolerant proteotypes were not uniform but formed two subclusters (Figure 5E). Differential expression identified 32 significant proteins (FDR ≤ 0.05, 26 up, 6 down; Figure 5F). The S100-high group was marked by elevated S100 family members and cornification-related proteins including CASP14, TGM1/TGM3, SBSN and DSC1, whereas the S100-low group showed higher levels of nuclear/chromatin related proteins (e.g., PTMA and HMGN4) and the oxidative-stress protector OXR1. Consistent with a less differentiated, expansion-competent state rather than increased S-phase engagement, S100-low colonies were significantly larger than S100-high colonies despite a similar EdU⁺ fraction (Figure 5G-I, Figure S5A). Together, these data suggest two phenotypically and proteomically distinct CP-tolerant states in primary HNSCC cells: an S100-high, cornification-biased state and an S100-low, chromatin-active state with greater colony expansion.

**Figure 5.**
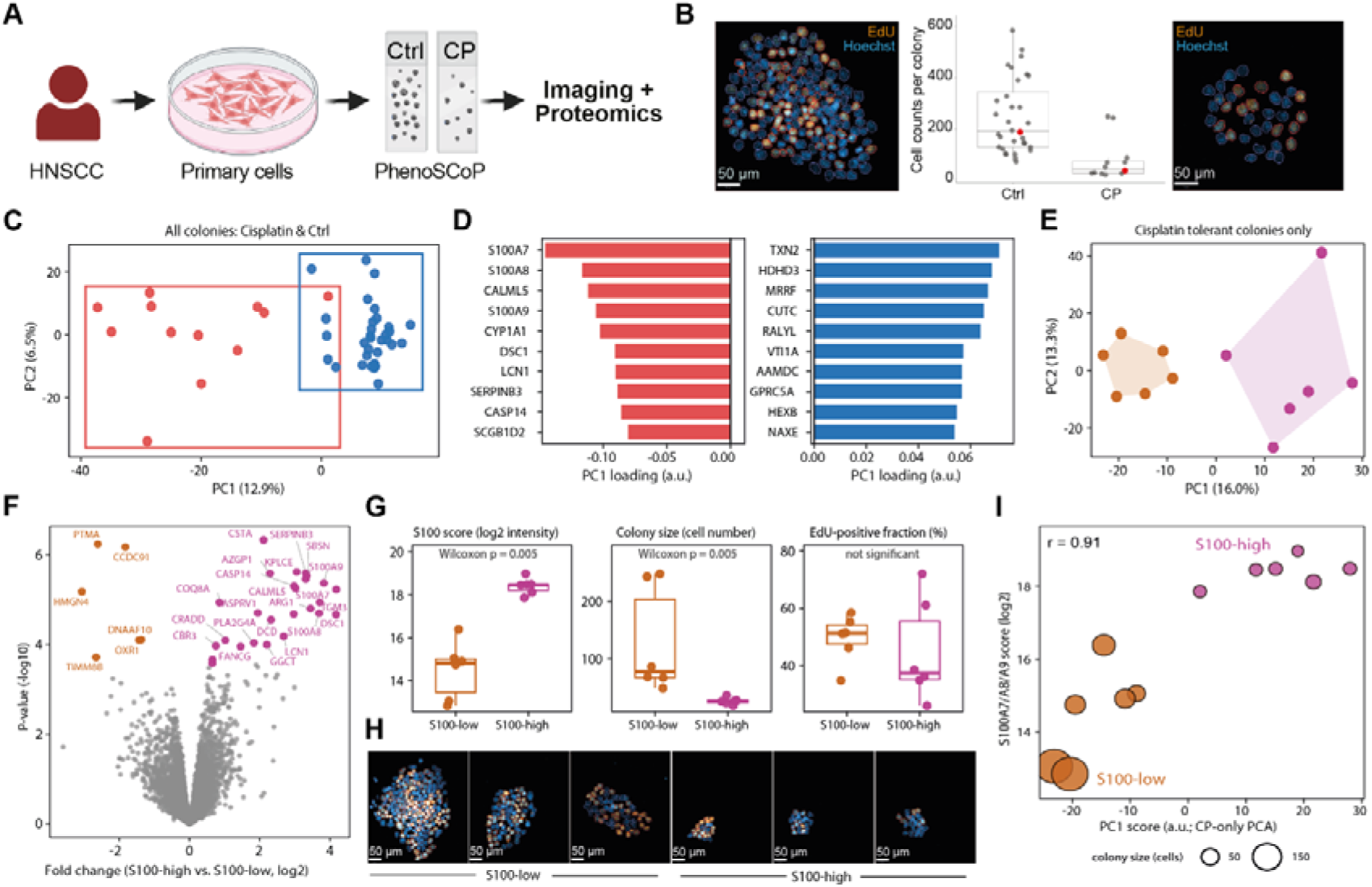
PhenoSCoP resolves heterogeneous CP-tolerant states in primary HNSCC cells. (A) Experimental workflow. Patient-derived primary HNSCC cells were seeded sparsely on membrane slides, treated with CP for 24 h or left untreated, and clonogenic outgrowth was monitored for 10 days. Individual colonies were imaged (including EdU labelling), laser microdissected, and analysed by single-colony proteomics. (B) Colony size (imaging-derived cell number) for untreated (Ctrl) versus CP-tolerant colonies. (C) Principal component analysis of all colony proteomes. Points are coloured by treatment. (D) Top PC1 loadings highlighting CP-associated cornification and squamous-differentiation proteins (e.g. S100A7/A8/A9, CALML5, SERPINB3, DSC1, CASP14). (E) PCA of CP-tolerant colonies coloured by S100A7/A8/A9 median-split subtype (S100-high versus S100-low), showing two proteomic subclusters. (F) Volcano plot of differential expression between S100-high and S100-low colonies. Proteins with FDR ≤ 0.05 are coloured (26 up, 6 down). The S100-high signature used for tissue analysis in Fig. 6 comprises the 26 upregulated proteins. (G) Comparison of the two subtypes: Left: S100A7/A8/A9 score, middle: colony size, right: EdU⁺ cell fraction. Boxplots show median and interquartile range; points are individual colonies. P values: two-sided Wilcoxon rank-sum tests. (H) Representative images of colonies from the S100-low and S100-high group. Blue: DAPI, orange: EdU. (I) Scatter plot of CP-tolerant primary HNSCC colonies (n = 12). The x-axis shows PC1 and the y-axis the S100A7/A8/A9 score (mean log₂ intensity). Points are coloured by median-split subtype (S100-high versus S100-low) and sized by colony size (cell number). Pearson correlation between S100 score and PC1 is indicated (r = 0.91, p = 3.5 × 10⁻⁵).

### PhenoSCoP-derived proteotypes are recapitulated in HNSCC patient tissues

Having resolved CP-tolerant S100-high and S100-low proteotypes in a primary HNSCC model (Figure 5), we next asked whether related programs could be identified in patient tissues. We first profiled a cohort of 29 HPV-negative, treatment-naïve HNSCC tumours from a previous phase III trial of definitive radiochemotherapy (RCTx) for locally advanced HNSCC (ARO-0401 cohort) ^44^, in which patients in the experimental arm received CP-based RCTx. Guided by H&E staining, tumour-specific compartments were laser-microdissected from tissue-microarray cores (three replicates per patient) and analysed by LC-MS (Figure 6A), yielding comparably deep tissue proteomes (Figure 6B, Table S6). Of the 357 variable proteins identified across FaDu colonies, 290 (81%) were detected in the ARO tissue cohort. Relative to the full tissue proteome, the FaDu-variable set of proteins showed significantly higher inter-patient variability (Figure 6C; P = 1.2 × 10⁻⁶). The most variable proteins included ALDH3A1, interferon-inducible proteins (e.g. ISG15, MX1 and IFIT2), GLUL and the calcium-binding proteins S100A8/A9 (Figure 6D).

**Figure 6.**
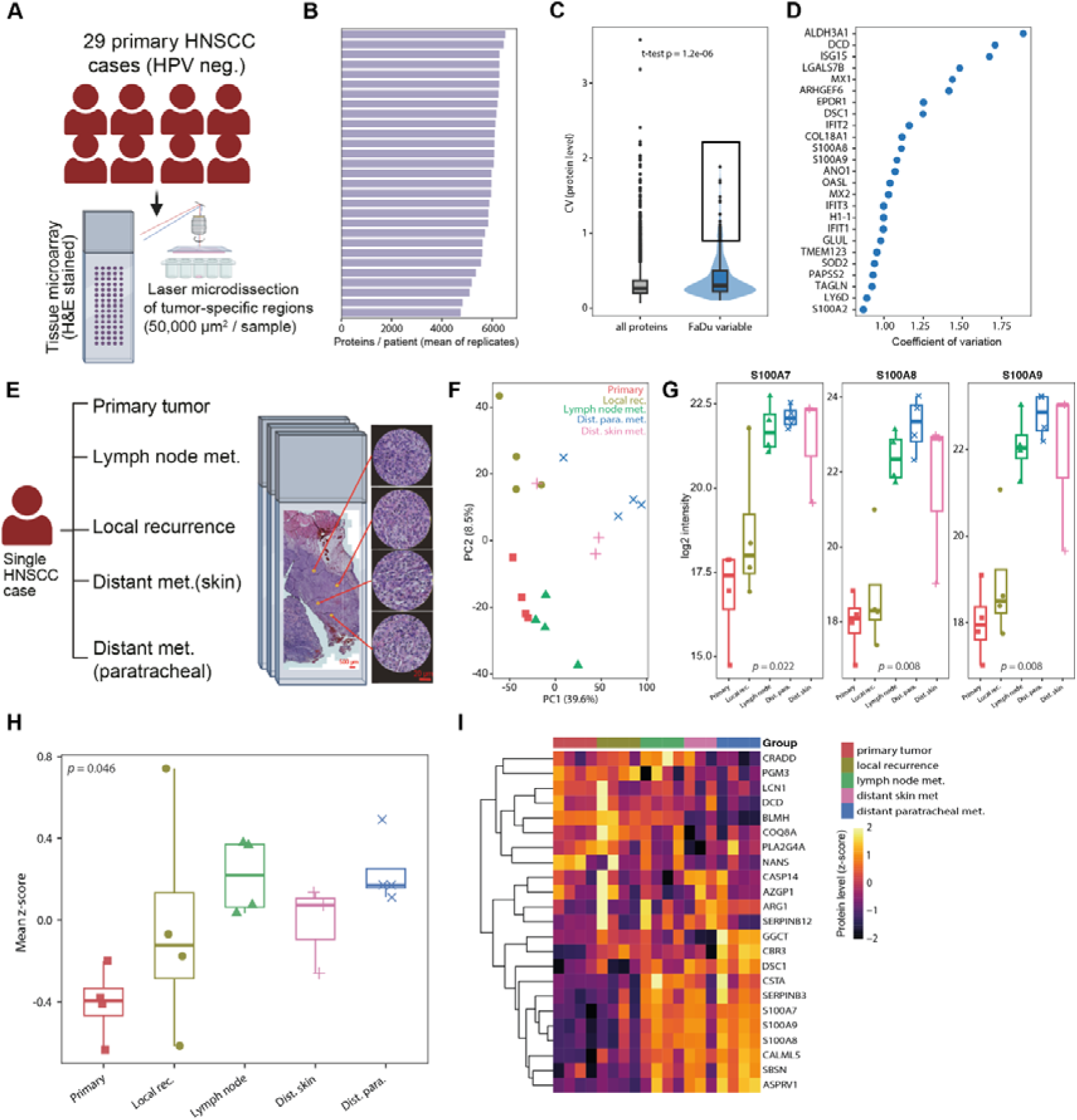
PhenoSCoP-derived proteotypes are recapitulated in HNSCC patient tissues. (A) Schematic workflow of laser microdissection guided proteomics applied to tissue microarrays of patients with primary head and neck squamous cell carcinoma (HNSCC). Tumor-specific compartments were identified by H&E staining, with three replicates per tissue cut for proteome analysis. (B) Protein identification in tissue samples from 29 patients with head and neck cancer. (C) Violin plot of inter-patient protein coefficients of variation comparing all quantified proteins to the variable proteins identified using PhenoSCoP in the FaDu cell line model. (D) Ranked inter-patient coefficients of variation for variable proteins (from the in vitro FaDu cell model). Proteins with the highest variability are listed and marked in blue. (E) Multi-site sampling of a CP-treated relapse case (primary tumour, local recurrence, lymph-node metastasis, distant paratracheal and skin metastases). (F) PCA of relapse microregion proteomes coloured by anatomic site; percentage variance explained by PC1 and PC2 is indicated. (G) Log₂ intensities of S100A7, S100A8 and S100A9 across sites; points are individual microregions. (H) Tissue proteomes scored using the primary-cell-derived S100-high signature (FDR ≤ 0.05 in Figure 5; 26 proteins; 23 detected in tissue). Signature score = mean z-scored abundance of signature proteins present in tissue. (I) Heatmap of S100-high signature proteins in tissue (z-scored). Columns are microregions ordered by anatomic site (colour bar).

To test whether these programs also track anatomic progression after CP exposure, we profiled a multi-site relapse case of an aggressive mandibular squamous cell carcinoma that recurred within six months after surgery with CP-based adjuvant RCTx and subsequently metastasized (Figure 6E). Microregions were sampled from primary tumour (n = 4), local recurrence (n = 4), lymph-node metastasis (n = 4), and distant metastases (paratracheal, n = 4; skin, n = 3), with comparable proteome depth across sites of ∼7,000 proteins per measurement (Figure S6A-B). Principal component analysis separated microregions by anatomic site, with distant metastases shifting along PC1 relative to primary/local disease (Figure 6F; PC1 39.6%, PC2 8.5%). Interestingly, one skin-metastasis and one paratracheal-metastasis microregion grouped closely with local-recurrence samples despite differing histomorphology in the corresponding H&E sections (Figure S6C). This finding indicates spatiotemporal proteomic relatedness among selected tumour regions that is not discernible by conventional morphological assessment, while the broader clustering pattern revealed substantial intra-patient proteomic heterogeneity.

Most proteins of the S100-high signature from primary cells were recovered in the tissue proteome (23/26). Consistent with the S100-high colony state, S100A7, S100A8 and S100A9 were lowest in primary tumour microregions and elevated in lymph-node and distant metastatic sites (Figure 6G; Kruskal–Wallis *p* = 0.022, 0.008 and 0.008, respectively). Scoring tissue samples with the colony S100-high signature (23 proteins) showed that S100-high scores differed across anatomic sites (Figure 6H; Kruskal–Wallis *p* = 0.046), with scores lowest in primary tumour and higher toward metastatic sites. Heatmap visualization of the colony-derived subtype signature proteins in tissue further separated primary/local from metastatic microregions (Figure 6I).

Together, these data indicate that PhenoSCoP-derived CP-tolerant proteotypes - particularly the S100-high cornification program - are detectable in HNSCC patient tissues and are enriched along a primary-to-metastatic anatomic axis.

## Discussion

Here, we introduce Phenotype-resolved Single-Colony Proteomics (PhenoSCoP), a versatile discovery proteomics approach that combines clonogenic outgrowth, fluorescence microscopy, laser microdissection, and ultrasensitive mass spectrometry-based proteomics. By profiling single-cell-derived colonies rather than isolated cells or bulk populations, PhenoSCoP achieves deep and reproducible proteome coverage while preserving clonal origin and image-defined phenotypes. Colony-level measurements reduce the influence of transient single-cell variation, including cell-cycle-associated effects, thereby prioritizing more persistent genetic and non-genetic clone-associated features. In this way, PhenoSCoP complements single-cell proteomics, which offers cellular resolution but typically loses clonal information and provides lower data completeness, and bulk proteomics, which averages out rare clonal states.

The FaDu model illustrates how this phenotype-to-proteome framework can uncover mechanisms of CP resistance. Resistant C5 and sensitive C46 clones displayed distinct proteomic programs that aligned with their genetic backgrounds. The resistant C5 clone showed stabilized mutant p53 together with elevated DNA-repair and replication-associated programs. Integration with colony imaging refined this interpretation: although C5 formed larger colonies and expressed higher levels of replication-associated proteins, it contained a lower fraction of EdU-positive cells at a given time. This suggests altered cell-cycle kinetics in which greater colony expansion coexists with reduced S-phase occupancy, potentially limiting the fraction of cells vulnerable to replication-associated cisplatin damage at any given time.

### PhenoSCoP therefore places proteomic signatures in the context of the proliferative composition of the clonal unit

Beyond genetically encoded differences, resistance can also arise through reversible, non-genetic adaptation. To capture these processes, we developed a live-cell colony-splitting strategy that provides matched proteomic measurements from the same clonally related population before and after CP exposure. By controlling for pre-existing inter-colony heterogeneity, this design distinguishes treatment-induced remodeling from stable clone-intrinsic proteotypes and extends endpoint proteomics toward temporal analysis of adaptation. NDUFB11, a mitochondrial complex I subunit, illustrates how this approach can prioritize functionally relevant vulnerabilities. NDUFB11 was downregulated after CP treatment, and its depletion preferentially impaired colony formation in the mutant p53-driven clone. Combined with CP, NDUFB11 depletion nearly abolished colony formation in the resistant clone. These findings suggest a clone-specific dependency on mitochondrial complex I function, although the underlying mechanism remains to be established.

PhenoSCoP also resolved pre-existing resistance-associated proteotypes within treatment-naïve parental populations. A small subset of untreated colonies already resembled resistant colonies generated by repeated CP exposure, a structure that mini-bulk measurements did not capture. RGB barcoding and image-guided selection further enable clonal tracking and prospective isolation of colonies with unusual phenotypic profiles.

Application of PhenoSCoP to patient-derived primary HNSCC cells further demonstrated that CP-tolerant colonies are molecularly and phenotypically heterogeneous. Stratification by an S100A7/A8/A9 score distinguished an S100-high state enriched for cornification and squamous-differentiation proteins (including CASP14, TGM1/TGM3, SBSN and DSC1) from an S100-low state characterized by nuclear and chromatin-associated proteins (including PTMA and HMGN4) and greater colony expansion, despite similar EdU-positive fractions. These findings suggest that differences in colony growth reflect distinct differentiation and expansion states. Several of these colony-derived programs were also observed in human HNSCC tissues. FaDu-variable proteins, including S100A8 and S100A9, were broadly detected in pretreatment tumors and showed elevated interpatient variability. Moreover, spatial profiling of one illustrative matched progression case showed increased S100A7/A8/A9 abundance from the primary tumor toward lymph-node and distant metastatic lesions. The convergence of S100-associated programs across cell-line colonies, patient-derived primary cells, and spatial tissue microregions suggests that epithelial differentiation and inflammatory stress programs may accompany therapy-tolerant and progression-associated states in HNSCC. Further functional studies will be required to determine whether S100A7/A8/A9 act as drivers of these phenotypes, biomarkers of aggressive cellular states, or both.

PhenoSCoP currently requires adherent cells capable of clonogenic outgrowth and is therefore less suitable for non-proliferative or poorly adherent states, which may be better addressed by direct single-cell microdissection approaches such as Deep Visual Proteomics ^58^. Colony-level averaging may also obscure rare or transient states within colonies. Although the current proteome coverage was sufficient to resolve major treatment-associated programs, further advances in analysis depth should be possible with newer instrumentation and optimized acquisition strategies.

In summary, PhenoSCoP bridges the gap between single-cell and bulk proteomics by linking quantitative proteome states to clonally resolved phenotypes and treatment histories. The approach distinguishes stable clone-intrinsic programs from adaptive responses, detects rare pre-resistant states obscured by population averaging, and prioritizes candidate vulnerabilities and biomarkers across cell lines, patient-derived primary cells, and tumor tissues. PhenoSCoP therefore provides a broadly applicable framework for investigating how clonal proteomic heterogeneity shapes therapeutic response and disease progression.

## Resource availability

### Lead contact

Further information and requests for resources and reagents should be directed to and will be fulfilled by the lead contact, Fabian Coscia.

### Materials availability

This study did not generate new materials.

## Data and code availability

- The mass spectrometry proteomics data have been deposited to the ProteomeXchange Consortium (http://proteomecentral.proteomexchange.org) via the PRIDE partner ^59^ with the dataset identifier PXD069541.
- This paper does not report original code.
- Any additional information required to reanalyse the data reported in this paper is available from the lead contact upon request.

## Author contributions

Conceptualization, D.Q. and F.C.; methodology, D.Q., S.S., R.D., K.R.; experiments, D.Q., R.D.; data curation, D.Q., S.S., I .T., F.C.; data analysis, D.Q. and F.C.; figures, D.Q., and F.C.; supervision, I .T. and F.C.; funding acquisition, K.R., I .T. and F.C.; writing the original draft, F.C. All authors reviewed and edited the manuscript.

## Acknowledgments

We thank our colleagues at the Max Delbrück Center (MDC) and Charité for their support and fruitful discussions. In particular, we thank Daniela Valdes and Bella Banjanin for their feedback on the manuscript. We also thank Gina Dörpholz, Pia Larsen and Jeannine Engel for administrative support. D.Q., F.C., I .T., and S.S. acknowledge funding support from the Federal Ministry of Education and Research (BMBF), as part of the National Research Initiatives for Mass Spectrometry in Systems Medicine, under grant agreement No. 161L0222 and 16LW0239K (Charité – University Medicine Berlin) of funding phase B, that is MSTARS-2. This project received funding from the European Research Council (ERC) under the European Union’s Horizon 2020 Research and Innovation Program (grant agreement No. 101115681) and was supported by the ERC (ERC starting grant). K.R. and I .T. were funded by the Deutsche Forschungsgemeinschaft (DFG, German Research Foundation) – Project number 467261817. Schematic workflow figures were created using BioRender.com.

## Declaration of generative AI and AI-assisted technologies in the writing process

During the preparation of this manuscript, the authors used Paperpal and ChatGPT in order to improve readability and language of the text. After using this tool, the authors reviewed and edited the content as needed and take full responsibility for the content of the publication.

## Methods

### HNSCC patient tissue collection

Primary cancer patient samples were obtained from a multicenter phase I II trial (ARO-0401) conducted between 2004 and 2008. All the samples included in this study (n = 29) were HPV-negative treatment-naïve tumors. Multiple tumor specimens from the recurrent patient (HPV-negative) were collected between 2022 and 2023 during surgery, including the primary tumor, lymph node metastasis, local recurrence, and distant metastases to the skin and paratracheal region. All tissue blocks were stored at room temperature in the archive of the Institute of Pathology at Charité University Hospital, Campus Benjamin Franklin. The study was performed according to the ethical principles for medical research of the Declaraûon of Helsinki, and approval was obtained from the Ethics Committee of the Charité University Medical Department in Berlin (EA1/222/21). Informed consent was obtained from all participants included in the study.

A patient-derived tumor organoid model, HN041, was included to evaluate the applicability of the established protocol to primary tumor cells. HN041 was established from a hypopharyngeal carcinoma specimen obtained during surgical resection under Institutional Review Board approval (EA2/152/10), with written informed consent from the patient. Organoid culture was established as previously described ^60^. The organoids were maintained under standard culture conditions (37°C, 5% CO₂) in complete patient-derived tumor organoid medium consisting of Advanced DMEM/F12 supplemented with N2, B27, EGF, bFGF, Y-27632, TGF-β receptor kinase inhibitor I V, Noggin, and Nutlin-3. Whole exome sequencing identified the TP53 p.C238S variant at a variant allele frequency of 100%, consistent with loss of the wild-type allele and confirming the predominantly tumor-derived nature of the culture. The HN041 model exhibited a cisplatin IC₅₀ of 1.78 µM, as determined using the CellTiter-Glo® cell viability assay.

### Cell lines

The hypopharyngeal tumor cell line FaDu (ATCC®HTB–43™) was purchased from ATCC (Manassas, VA, USA). FaDu clonal cell lines (C5 and C46) were generated in our previous study ^43^. The cells were cultured in Minimum Essential Medium (MEM) supplemented with 12% fetal bovine serum (FBS), 1× non-essential amino acids (NEAA), and penicillin/streptomycin. Cells were regularly tested for mycoplasma by PCR.

### Transduction of FaDu cells with RGB

FaDu cells were transduced with the lentiviral vectors LeGO-B2-NLS-Puro+, LeGO-V2-NLS-Puro+, and LeGO-C2-NLS-Puro+ for RGB nuclear barcoding, followed by molecular barcoding with LeGO-EBFP2-Hygro-LTRXX3-BC24 for clonal tracking. For transduction, 50,000 cells per well were seeded in a 24-well plate (Greiner Bio-One #353047). Cells were incubated for 2-5 hours to facilitate cell attachment. The medium was replaced with polybrene-containing medium (8 μg/mL), and lentiviral particles were added to the wells. Spinoculation (1,000 × g, 1 h, 25 °C) was performed on the plate, and cells were incubated for another 16 h. The medium was then exchanged for a regular, non-polybrene-containing medium.

The efficiency of RGB transduction was determined by flow cytometry. The most appropriate transduction combination (all three 3 vectors) for maximal color distribution was used for molecular barcoding. The efficiency of molecular barcoding was determined by ddPCR using the ratio of Hygro/FAM9B. A ratio of 1 or the closest to 1 was used for further experiments with irradiation (IRR) and drug treatment.

RGB fluorescence imaging during colony seeding was used as a quality-control step to verify colony integrity before laser microdissection. Colonies exhibiting obvious mixtures of fluorescent labels, indicative of colony overlap or fusion between independently labeled cells, were excluded from further analysis. Relative RGB protein abundances were quantified by mass spectrometry using a customized FASTA database containing the RGB fluorescent protein sequences.

### Colony growth for PhenoSCoP

Steel-frame PPS (polyphenylene sulfide) membrane slides (Leica, #11600294) and 12 mm round glass coverslips were sterilized by exposure to ultraviolet (UV) light for 30 min before use. They were then incubated with 0.1 mg/mL poly-L-lysine (Sigma #P1274) at 37°C for 30 min, rinsed once with sterilized water, and allowed to dry under UV light. For seeding, 4,000–5,000 cells were placed into a 10 cm cell culture dish containing a single PPS slide, while approximately 500 cells were seeded into each well of a 24-well plate, each containing a sterilized coverslip. Cells were seeded on the upward-facing PPS membrane surface, preventing growth underneath the slide. The culture medium was replaced every 2–3 days to ensure optimal cell growth. Colonies located at the interface between the PPS membrane and the metal frame, which could not be completely isolated by laser microdissection, were excluded from analysis. Only colonies entirely contained within the PPS membrane area were selected for proteomic profiling.

### Drug treatment

Cells were seeded onto membrane slides and incubated overnight at 37°C to allow cell attachment. The following day, cells were treated with cisplatin at concentrations corresponding to their individual IC₅₀ values for 24 h. After treatment, the cells were washed once with phosphate-buffered saline (PBS), and fresh drug-free medium was added. The medium was replaced at regular intervals to support colony growth.

For cyclic CP treatment, 30,000 FaDu cells were seeded in each well of a 6-well plate. Cells were treated with four doses of 100 ng/mL and four doses of 150 ng/mL, with a cumulative dose of 1000 ng/ml. Cells were split and reseeded after every cisplatin dose and harvested for further experiments. The cisplatin used for drug testing is from NeoCorp GmbH, Germany.

### SiRNA transfection

siRNA pools (siPOOLs) were obtained from siTOOLs Biotech GmbH. Gene knockdown was performed in untreated FaDu parental cells and the cisplatin-resistant clone (C5) using either control siRNA (siCTRL) or an siPOOL targeting NDUFB11, according to the manufacturer’s instructions. Briefly, 240,000 cells were seeded per well in 6-well plates and allowed to adhere overnight prior to transfection. Cells were transfected with 3 nM siRNA and incubated for 48 h. For the clonogenic assay, 500 cells were seeded per condition.

### Immunofluorescence microscopy

After 8–10 days of growth, colonies were fixed with 4% paraformaldehyde (PFA, Thermo Scientific, # 28908) for 10 min at room temperature. A silicon frame (0.5 mm thickness, Technikplaza GmbH) was mounted onto the slide to facilitate the subsequent steps. Immunofluorescence staining was performed on PPS slides. In brief, cells and colonies were permeabilized by adding 90% cold methanol (in 10% PBS) for 2 min. After thoroughly washing away the methanol, Odyssey Blocking Buffer PBS (Li-Cor, 927-40000) was applied to block non-specific binding sites for 30 min at room temperature. Antibodies were diluted to optimal concentrations in blocking buffer and incubated with the cells overnight at 4°C. When unconjugated primary antibodies were used, a secondary antibody was added and incubated for 1 h at room temperature. Hoechst 33342 (Thermo Scientific, # 62249) was applied for 10 min at room temperature for nuclear staining. Images were acquired using an Axioscan 7 (Carl Zeiss Microscopy GmbH, Germany) with a 10x objective. The optimal antibody concentrations used were as follows: Na/K ATPase (1:500, Abcam, # 76020), panCK (1:100, Thermo Fisher Scientific, 41-9003-82), DDX60 (1:20, Novus, NBP1-91826), TP53 (1:50, Agilent Dako, M7001), and Ki-67 (1:100, Cell Signaling Technology, #11882). The secondary antibodies used were goat anti-mouse (1:500, Alexa Fluor™ 750, Thermo Fisher Scientific, A-21037) and goat anti-rabbit (1:1000, Alexa Fluor™ 488, Thermo Fisher Scientific). EdU staining was performed using a commercial kit according to the manufacturer’s instructions (Thermo Fisher Scientific, C10338).

### Laser microdissection and proteomic sample preparation

Colony selection was performed using QuPath (v0.5.1) and exported to LMD using an in-lab pipeline (https://github.com/CosciaLab/Qupath_to_LMD). Laser microdissection was performed on an LMD7 instrument (Leica Microsystems, Wetzlar, Germany) using a 5× objective. The collection efficiency exceeded 98%, with nearly all dissected colony regions successfully transferred into their designated wells. Sample preparation for colony and patient samples was performed based on our previous work ^61^. Briefly, samples in 384-well plates were lysed with 2 µL of lysis buffer consisting of 0.1% DDM, 5 mM TCEP, 20 mM CAA, and 100 mM TEAB at 95 °C for 1 h. Sequential digestion was applied by adding 1 µL of LysC (2 ng) and incubating for 4 h, followed by the addition of 1 µL of trypsin (2 ng) and overnight incubation at 37 °C. All buffers were added using a liquid dispenser (MANTIS, Formulatrix) for better accuracy. Peptides were desalted using Evotips (Evosep Biosystems, Denmark) according to the manufacturer’s instructions. Eluted peptides were dried using a SpeedVac and resuspended in 4.2 µL of buffer A* (3% ACN, 0.1% FA). Injection volumes were adjusted using image-derived estimates of cellular input. Thus, injection volume was calculated as: injection volume = total digest volume × target cell equivalent / estimated colony cell number. Patient-derived primary-cell colonies were normalized to 50-cell equivalents using the relationship between colony area and image-derived cell number.

For live-colony splitting experiments, regions of approximately 80,000–100,000 µm² were isolated from the peripheral edge of each colony using a 10× objective. Colonies were maintained in 1 mL of prewarmed phosphate-buffered saline (PBS), and the cutting chamber was maintained at 37°C to preserve cell viability. The total processing time for each slide was limited to 1 h. Laser microdissection was performed in two sequential steps: cutting and extraction. During the cutting step, the instrument was operated in Draw + cut mode (laser power 40, aperture 1, speed 4), and each region was cut twice to ensure complete separation. During the extraction step, Move-and-cut mode was used (laser power 60, aperture 27, speed 4). Dissected colony fragments were collected into a 384-well plate and transferred to the bottom of each well by rinsing with acetonitrile for downstream proteomic sample preparation, with gentle manipulation using a pipette tip when necessary. Each well was inspected microscopically after collection to confirm successful transfer. Samples collected before treatment were stored at −20°C and processed in parallel with samples collected after treatment, as described above.

### Single-cell proteomics

Single FaDu cells were sorted and prepared using the CellenONE system (Cellenion) on proteoCHIP EVO 96 plates according to an adapted published protocol ^62^. Briefly, 3 µL of hexadecane was added to each well prior to cell sorting, followed by dispensing 150 nL of Master Mix (0.2% DDM, 100 mM TEAB, and 10 ng/µL trypsin) into each well. During lysis and enzymatic digestion, the on-deck temperature was maintained at 50 °C, with continuous rehydration of samples by adding 60 nL H₂O at 2-minute intervals for 2 h. Subsequently, the samples were loaded onto Evotips by centrifugation.

### Liquid chromatography-mass spectrometry (LC-MS/MS)

LC-MS/MS was performed using timsTOF Ultra/Ultra 2 mass spectrometers (Bruker Daltonics, Germany) coupled to an EASYnLC-1200 system (Thermo Fisher Scientific, USA). Peptide separation was conducted on 20 cm home-packed columns (75 μm inner diameter) packed with 1.9 μm ReproSil-Pur C18-AQ silica beads (Dr. Maisch GmbH, Germany). A 21-minute total gradient (15-min active gradient) was used for peptide separation, employing buffer A (3% acetonitrile [ACN], 0.1% formic acid) and buffer B (90% ACN, 0.1% formic acid). The gradient was initiated at 2% buffer B and increased to 10% over 1.5 minutes at a flow rate of 0.4 μL/min to minimize dead time at the start of chromatography. Subsequently, the flow rate was reduced to 0.25 μL/min, and buffer B was ramped up to 60% over 15 min, followed by a washing step at 90% buffer B. All mass spectrometric analyses were conducted using the dia-PASEF method ^33^, employing the factory default settings in high sensitivity mode. Eight dia-PASEF windows were distributed across three trapped ion mobility spectrometry (TIMS) scans, covering an m/z range of 400 to 1,000. The ion mobility range was set from 0.64 to 1.37 Vs cm ⁻ ², with an accumulation and ramp time of 100 ms.

Primary patient samples were measured using the Evosep ONE 30 SPD system with a 15 cm column (EV1106). Multi-cancer samples from the recurrent patient and single-cell samples were analyzed using the Whisper Zoom 40 SPD system (Evosep) equipped with an Aurora Elite column (15 × 75 µm, 1.7 µm).

### Raw file analysis

We used DIA-NN ^34^ (versions 1.8.1 and 1.9) for proteome quantification. Colony and patient samples were searched against an in silico spectral library generated from the UniProt human reference proteome (2023 release) supplemented with a FASTA file containing common contaminants. In addition, three color vector sequences (RGB) were included in the library to enable RGB sample identification. For single-cell samples, a FaDu cell–specific spectral library was generated using eight fractions of pooled cell lysate. Carbamidomethylation of cysteine residues was specified as a fixed modification, whereas oxidation of methionine was included as a variable modification. The maximum number of missed cleavages and variable modifications per peptide was set to one. Match-between-runs (MBR) functionality was enabled for all analyses.

### Data analysis

Data analysis was carried out using the pg.matrix output file of DIA-NN with a global protein FDR of max. 1%. Perseus (v1.6.15.0) and R (v4.4.1) were used for further data analysis. The following R packages were used: tidyverse (2.0.0), ggplot2 (4.0.0), dplyr (1.1.4), cluster (2.1.8.1), factoextra (1.0.7), stringr (1.5.1), tidyr (1.3.1), clusterProfiler (4.15.1.2), doBy (4.7.0), ComplexHeatmap (2.20.0), circlize (0.4.16).

Variable proteins in primary FaDu cells were calculated as follows: The data were filtered to retain proteins with at least five valid replicates. The intensity values were then log2-transformed to prepare for the imputation of missing values. Missing values were replaced by sampling from a normal distribution. The coefficient of variation (CV) was calculated in linear space, while the mean expression level for each protein was log-transformed. To obtain variable proteins across the whole dynamic range, the mean expression levels of proteins across colonies were divided into 100 bins. The top 5% of proteins with the highest CV were selected from each bin. Low-intensity proteins below 5% of the colony mean distribution were removed to reduce the influence of technical noise on the results. Proteins were excluded if their CV in mini-bulk samples was larger than that of the colony samples.

## Supplementary Figures

**Figure S1.**
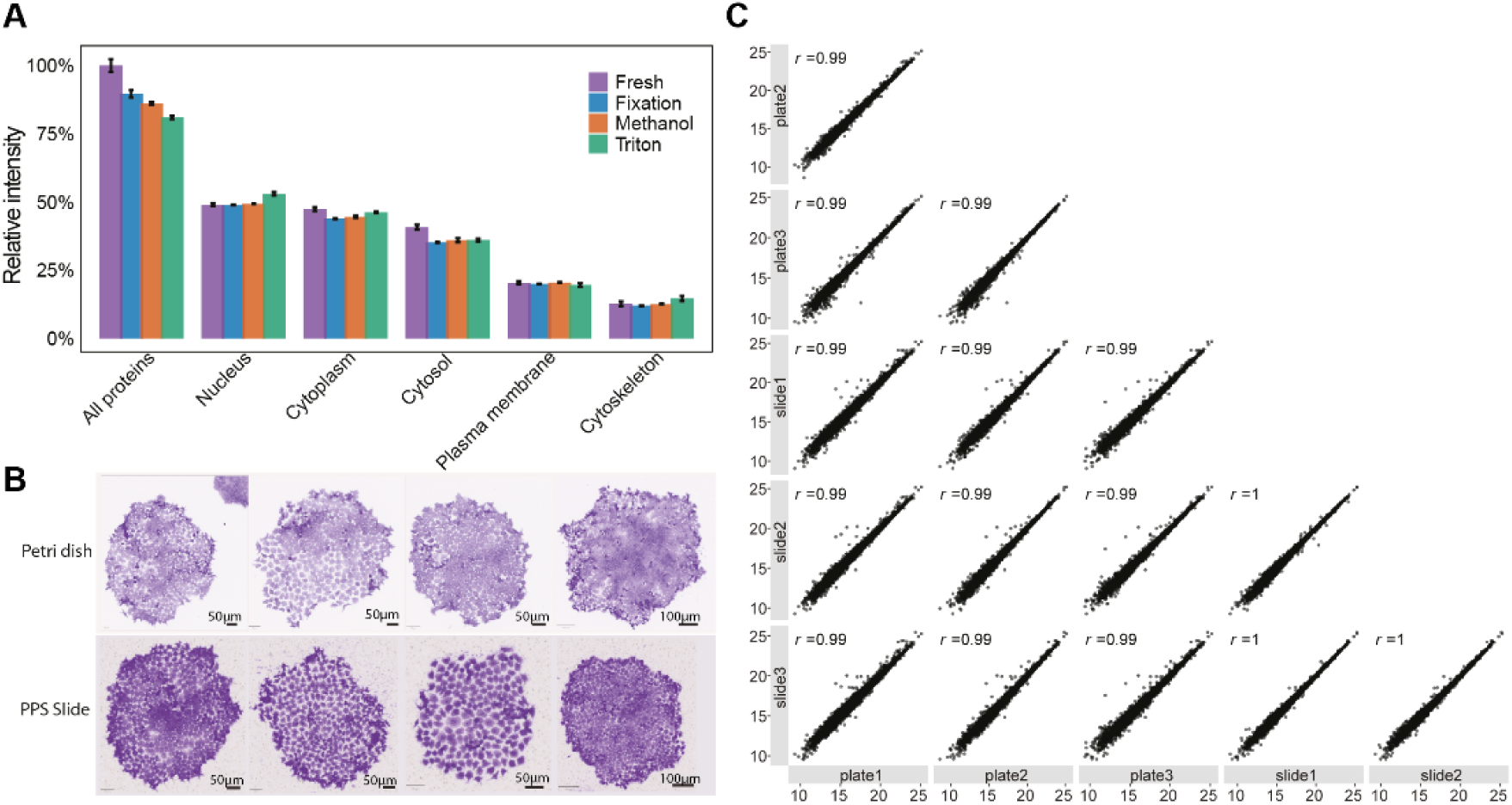
Impact of fixation and permeabilization on the proteomic read-out. (A) Comparison of summed intensity ratios from different cellular compartments across four experimental conditions: Fresh cells, formaldehyde fixed cells (FA, fixation), FA fixed cells and permeabilized cells based on ice-cold methanol (Methanol) and Triton-X-100 (Triton). ‘Sum’ represents the total intensity of proteins shared across all conditions (normalized to the average intensity of fresh cells). Other cellular compartments represent the summed intensity of proteins assigned to Gene Ontology Cellular Component groups (normalized to the total protein intensity of each sample). (B) Representative images of crystal violet-stained colonies cultured on conventional plastic dishes or PPS membrane slides. Images were acquired using a Leica MICA microscope at 1.6× magnification. (C) Pairwise correlation of proteomic profiles from cells digested from conventional plastic dishes (plates) and PPS membrane slides. Three technical replicates were analyzed for each culture condition. Pearson correlation coefficients (R) are shown for each pairwise comparison.

**Figure S2.**
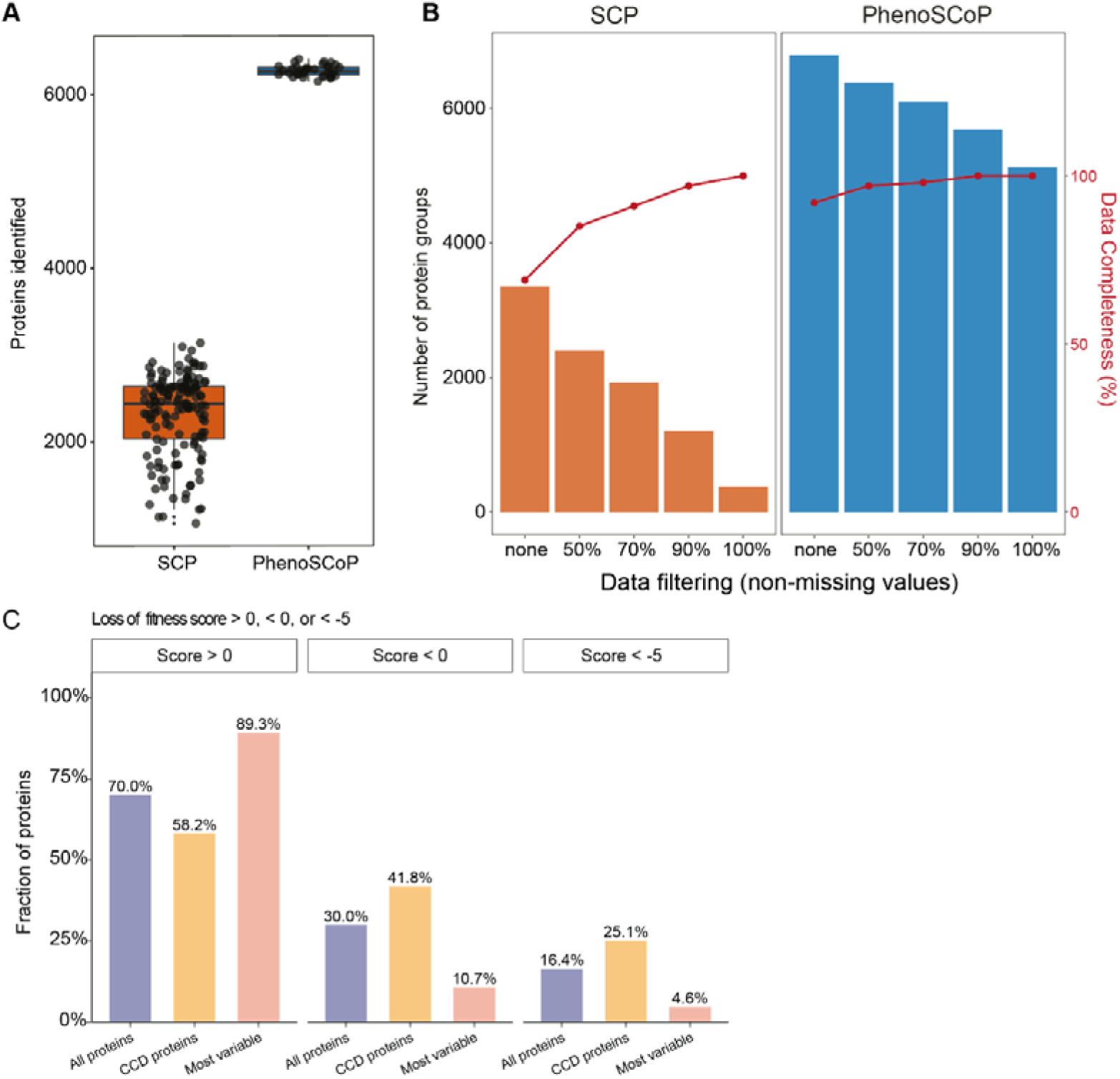
Comparison of proteome depth, data completeness, and protein essentiality between single-cell proteomics and PhenoSCoP. (A) Boxplot showing the number of proteins quantified using single-cell proteomics (SCP; n = 147) and PhenoSCoP (n = 36). (B) Comparison of protein group numbers and data completeness under different data-filtering criteria in SCP and PhenoSCoP. ‘None’ indicates no data filtering. (C) FaDu gene dependency scores (loss of fitness score; more negative = stronger essentiality) for proteins quantified in colony proteomes and detected in ≥50% of colony samples (n = 6,236). Proteins were grouped as all proteins, Human Protein Atlas cell cycle–dependent (CCD) proteins (n = 323), or the most variable colony proteins (top 10% coefficient of variation, CV; n = 624). Shown is the fraction of proteins with score > 0, < 0, or < −5 per category.

**Figure S3.**
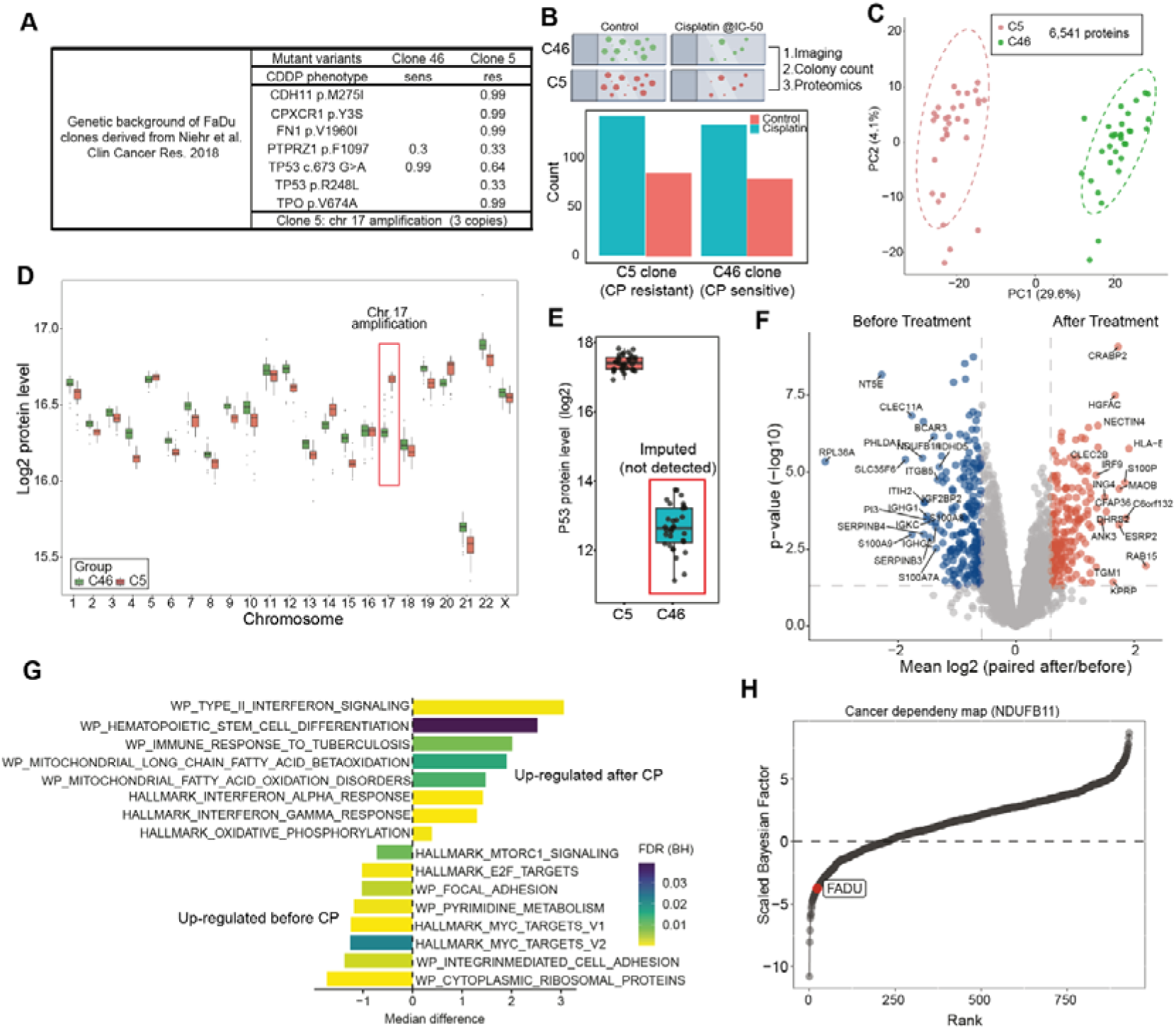
Stable clone-intrinsic programs and treatment-induced adaptations in CP resistance. (A) Genetic background of clone-derived cell lines from FaDu. (B) Top: Representative images of untreated and CP-treated (IC₅₀) colonies grown on membrane slides. Bottom: Quantification of colony numbers for CP-sensitive (C46, IC_50_: 30 ng/mL) and CP-resistant (C5, IC_50_: 150 ng/mL) clones under untreated and respective IC₅₀ treatment conditions. (C) Principal component analysis (PCA) of C5 (n = 32) and C46 (n = 32) colonies analyzed using PhenoSCoP. (D) Comparison of protein levels by chromosomal location. Note that C5 carried chromosome 17 amplification. (E) Boxplot of p53 protein level (Log2) in C5 and C46 colonies. (F) Volcano plot showing differential protein abundance between paired colonies before and after CP treatment (n = 18 paired colonies). Protein abundance was compared within each colony pair using paired statistical analysis. (G) Pathway enrichment analysis of paired colonies before and after CP exposure, using WikiPathways and Hallmark gene sets. (H) Ranked plot of scaled Bayes factors for NDUFB11 dependency across 930 cancer models. The score of the FaDu cell line is indicated in the plot. (CRISPR: Dependencies data were downloaded from the Sanger Institute).

**Figure S4.**
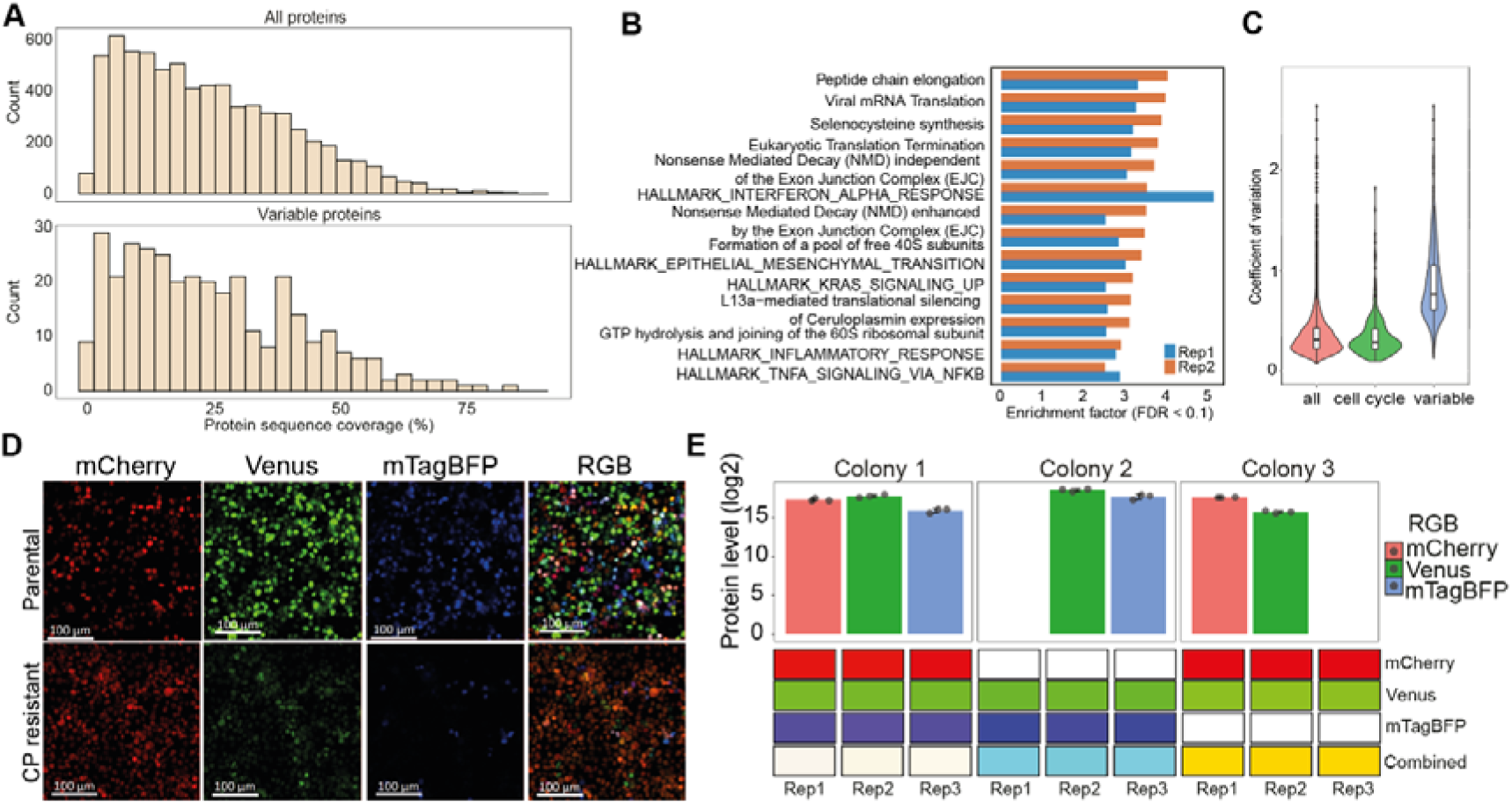
Variable protein programs and RGB barcoding across FaDu colonies. (A) Histogram of proteomic coverage for all proteins and variable proteins. (B) Pathway enrichment analysis of variable proteins in repeat experiments. The HALLMARK gene sets were obtained from the Human Molecular Signatures Database (MSigDB) and Reactome pathways. (C) Violin plots showing the distribution of the coefficient of variation (CV) for all proteins, cell cycle-related proteins (Mahdessian et al., 2021b), and variable proteins. (D) Representative images of RGB-labelled primary and resistant FaDu cells. Individual fluorescence channels are shown: red (mCherry), green (Venus), and blue (mTagBFP), as well as the merged RGB image. Scale bar: 100 µm. (E) Quantified RGB color values for three spatial subsamples from each of three colonies (n = 9 measurements). Upper panel: Bar plots display the mean log₂-transformed protein levels corresponding to the red, green, and blue components, with error bars representing the standard deviation across replicates. The protein abundance for each RGB vector was normalized to a 0-255 scale across all colonies. Lower panel: RGB values for individual replicates are shown separately for each colony along with the corresponding composite virtual RGB colors generated from the quantified levels.

**Figure S5.**
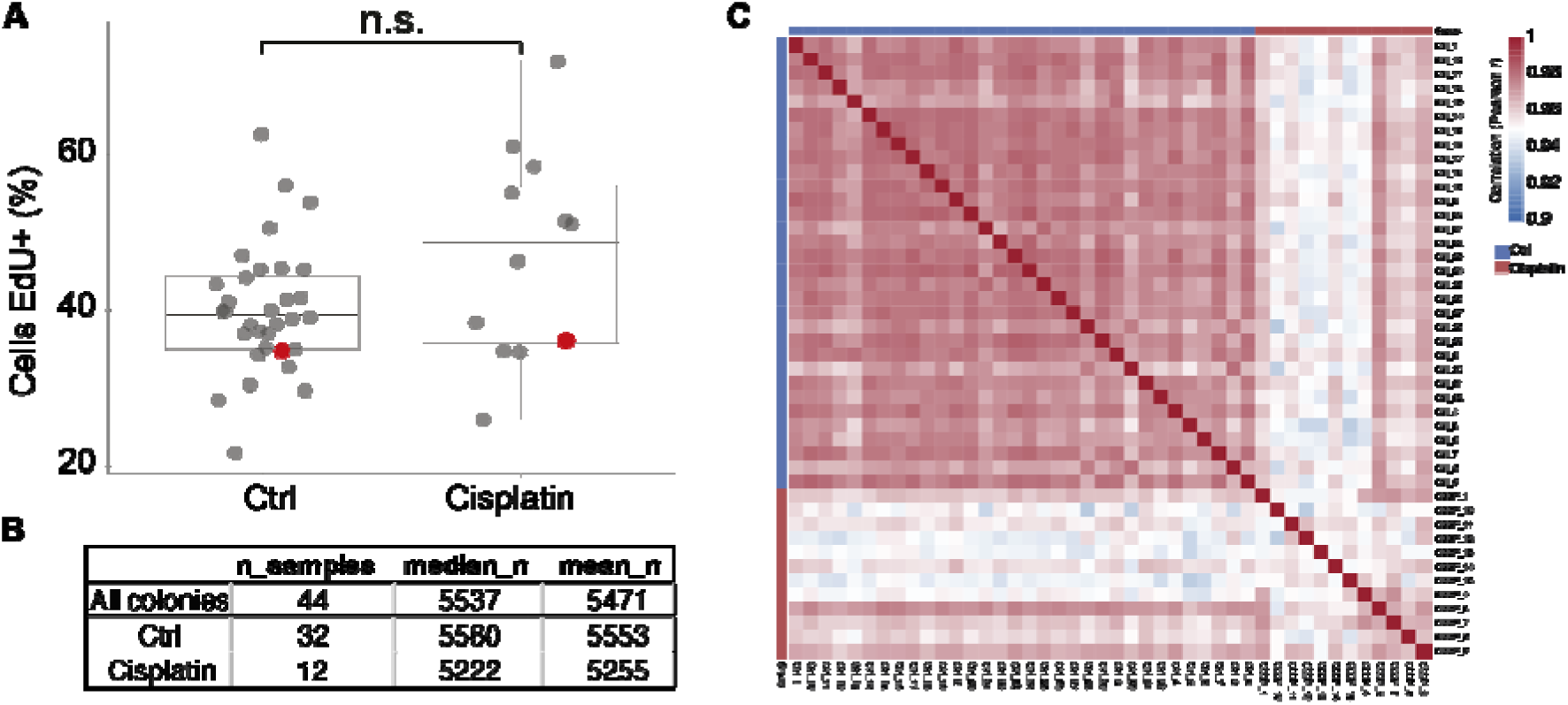
Quality control and extended analyses for primary HNSCC colony proteomics. (A) EdU labelling of CP-tolerant colonies after drug washout, showing recovery of proliferative capacity. Highlighted colonies in red are shown in Figure 5B. (B) Proteome coverage per colony from unimputed data (median 5,537). (C) Sample-sample Pearson correlation matrix of unimputed colony proteomes (Ctrl vs CP; median r = 0.97).

**Figure S6.**
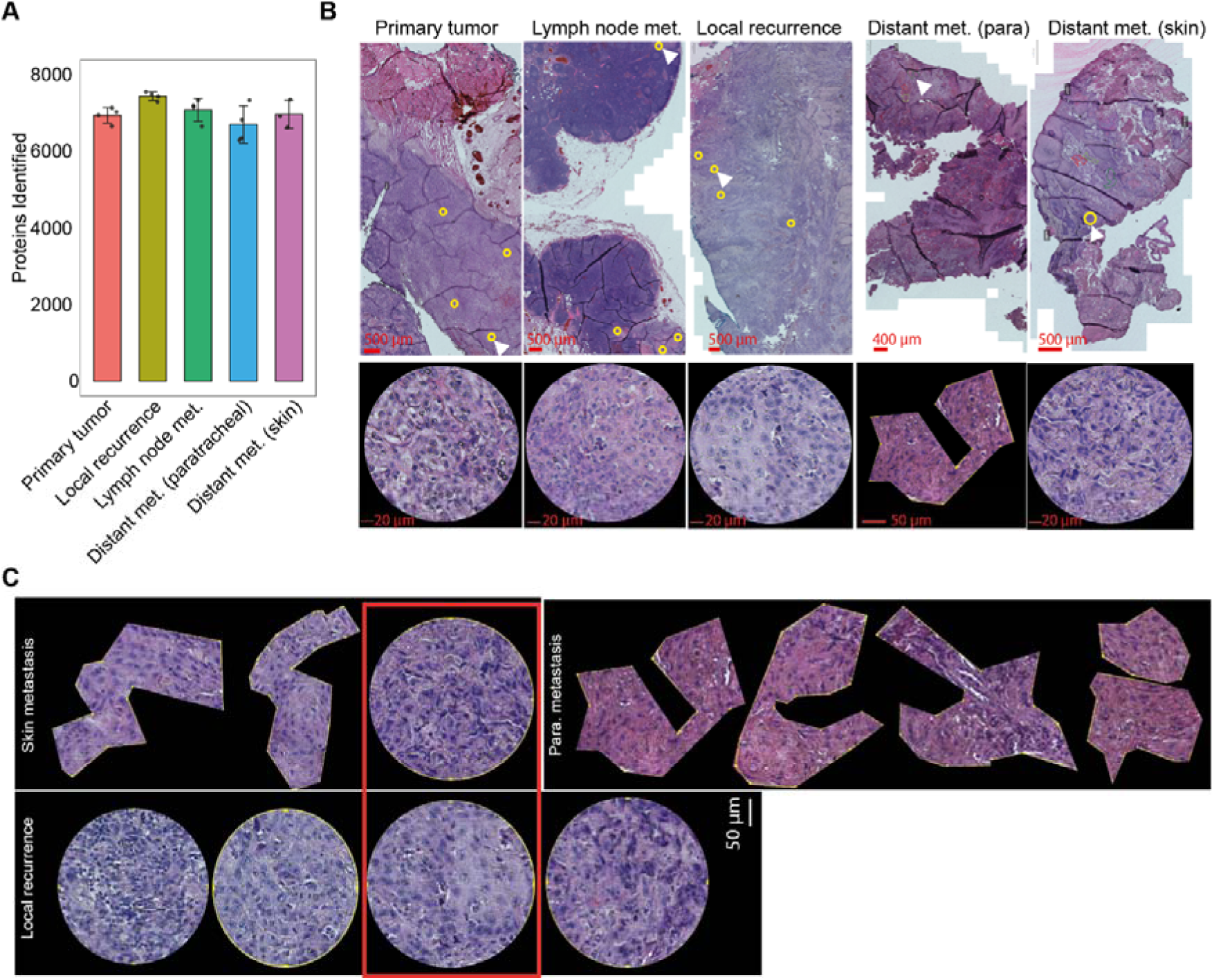
Spatial proteomics links colony-derived variability signatures to human HNSCC tissues. (A) Bar chart showing the number of proteins identified in multiple cancer tissue samples from a patient with relapse, including the primary tumor (n = 4), lymph node metastasis (n = 4), local recurrence (n = 4), and two distant metastases (paratracheal [n = 4] and skin [n = 3]). (B) Representative H&E images of multiple cancer tissue samples from a patient with relapse, including the primary tumor, lymph node metastasis, local recurrence, and two distant metastases (paratracheal and skin). Example regions isolated for proteomic measurements are shown in the lower panel. (C) Representative images of the sampled regions from the local recurrence, skin and paratracheal metastasis specimens. Regions of interest (ROIs) from the two specimens that clustered together in the principal component analysis (PCA) are outlined.

## References

1. Vasan, N., Baselga, J., and Hyman, D.M. (2019). A view on drug resistance in cancer. Nature 575, 299–309. 10.1038/s41586-019-1730-1.

2. Dagogo-jack, I., and Shaw, A.T. (2017). Tumour heterogeneity and resistance to cancer therapies. Nature Publishing Group 15, 81–94. 10.1038/nrclinonc.2017.166.

3. Mcgranahan, N., and Swanton, C. (2017). Clonal Heterogeneity and Tumor Evolution: Past, Present, and the Future. Cell 168, 613–628. 10.1016/j.cell.2017.01.018.

4. Marine, J.C., Dawson, S.J., and Dawson, M.A. (2020). Non-genetic mechanisms of therapeutic resistance in cancer. Nature Reviews Cancer 2020 20:12 20, 743–756. 10.1038/s41568-020-00302-4.

5. Nam, A.S., Chaligne, R., and Landau, D.A. (2021). Integrating genetic and non-genetic determinants of cancer evolution by single-cell multi-omics. Nat. Rev. Genet. 22, 3–18. 10.1038/s41576-020-0265-5.

6. Aebersold, R., and Mann, M. (2016). Mass-spectrometric exploration of proteome structure and function. Nature 537, 347–355. 10.1038/nature19949.

7. Goyal, Y., Busch, G.T., Pillai, M., Li, J., Boe, R.H., Grody, E.I ., Chelvanambi, M., Dardani, I .P., Emert, B., Bodkin, N., et al. (2023). Diverse clonal fates emerge upon drug treatment of homogeneous cancer cells. Nature 620, 651–659. 10.1038/s41586-023-06342-8.

8. Schaff, D.L., Fasse, A.J., White, P.E., Vander Velde, R.J., and Shaffer, S.M. (2024). Clonal differences underlie variable responses to sequential and prolonged treatment. Cell Syst. 15, 213–226.e9. 10.1016/j.cels.2024.01.011.

9. Sydney Shaffer, A.M., Emert, B.L., Reyes Hueros, R.A., Singh, A., Bassett, D.S., and Raj Correspondence, A. (2020). Memory Sequencing Reveals Heritable Single-Cell Gene Expression Programs Associated with Distinct Cellular Behaviors. 10.1016/j.cell.2020.07.003.

10. Rambow, F., Rogiers, A., Marin-Bejar, O., Aibar, S., Femel, J., Dewaele, M., Karras, P., Brown, D., Chang, Y.H., Debiec-Rychter, M., et al. (2018). Toward Minimal Residual Disease-Directed Therapy in Melanoma. Cell 174, 843–855.e19. 10.1016/j.cell.2018.06.025.

11. Kim, C., Gao, R., Sei, E., Brandt, R., Hartman, J., Hatschek, T., Crosetto, N., Foukakis, T., and Navin, N.E. (2018). Chemoresistance Evolution in Triple-Negative Breast Cancer Delineated by Single-Cell Sequencing. Cell 173, 879–893.e13. 10.1016/J.CELL.2018.03.041.

12. Mahdessian, D., Cesnik, A.J., Gnann, C., Danielsson, F., Stenström, L., Arif, M., Zhang, C., Le, T., Johansson, F., Shutten, R., et al. (2021). Spatiotemporal dissection of the cell cycle with single-cell proteogenomics. Nature 590, 649–654. 10.1038/s41586-021-03232-9.

13. Cohen, A.A., Geva-Zatorsky, N., Eden, E., Frenkel-Morgenstern, M., Issaeva, I ., Sigal, A., Milo, R., Cohen-Saidon, C., Liron, Y., Kam, Z., et al. (2008). Dynamic proteomics of individual cancer cells in response to a drug. Science (1979). **322**, 1511–1516. 10.1126/science.1160165.

14. Snijder, B., Vladimer, G.I., Krall, N., Miura, K., Schmolke, A.S., Kornauth, C., Lopez de la Fuente, O., Choi, H.S., van der Kouwe, E., Gültekin, S., et al. (2017). Image-based ex-vivo drug screening for patients with aggressive haematological malignancies: interim results from a single-arm, open-label, pilot study. Lancet Haematol. 4, e595–e606. 10.1016/S2352-3026(17)30208-9.

15. Lin, J.R., Fallahi-Sichani, M., and Sorger, P.K. (2015). Highly multiplexed imaging of single cells using a high-throughput cyclic immunofluorescence method. Nature Communications 2015 6:1 **6**, 1–7. 10.1038/ncomms9390.

16. Mills, C.E., Subramanian, K., Hafner, M., Niepel, M., Gerosa, L., Chung, M., Victor, C., Gaudio, B., Yapp, C., Nirmal, A.J., et al. (2022). Multiplexed and reproducible high content screening of live and fixed cells using Dye Drop. Nat. Commun. 13, 1–18. 10.1038/s41467-022-34536-7.

17. Bennett, H.M., Stephenson, W., and Rose, C.M. (2023). Single-cell proteomics enabled by next-generation sequencing or mass spectrometry. Nat. Methods. 10.1038/s41592-023-01791-5.

18. Geiger, T. (2023). Tackling tumor complexity with single-cell proteomics. 20, 324–326. 10.1038/s41592-023-01784-4.

19. Leduc, A., Huffman, R.G., Cantlon, J., Khan, S., and Slavov, N. (2022). Exploring functional protein covariation across single cells using nPOP. Genome Biol. 23, 1–31. 10.1186/s13059-022-02817-5.

20. Ye, Z., Sabatier, P., van der Hoeven, L., Lechner, M.Y., Phlairaharn, T., Guzman, U.H., Liu, Z., Huang, H., Huang, M., Li, X., et al. (2025). Enhanced sensitivity and scalability with a Chip-Tip workflow enables deep single-cell proteomics. Nat. Methods. 10.1038/s41592-024-02558-2.

21. Brunner, A., Thielert, M., Vasilopoulou, C., Ammar, C., Coscia, F., Mund, A., Hoerning, O.B., Bache, N., Apalategui, A., Lubeck, M., et al. (2022). Ultra-high sensitivity mass spectrometry quantifies single-cell proteome changes upon perturbation. Mol. Syst. Biol. 18. 10.15252/msb.202110798.

22. Gatto, L., Aebersold, R., Cox, J., Demichev, V., Derks, J., Emmott, E., Franks, A.M., Ivanov, A.R., Kelly, R.T., Khoury, L., et al. (2022). Initial recommendations for performing, benchmarking, and reporting single-cell proteomics experiments. 20, 375–386. 10.1038/s41592-023-01785-3.

23. Woo, J., Loycano, M., Amanullah, M., Qian, J., Amend, S.R., Pienta, K.J., and Zhang, H. (2025). Single-Cell Proteomic Characterization of Drug-Resistant Prostate Cancer Cells Reveals Molecular Signatures Associated with Morphological Changes. Molecular & Cellular Proteomics 24, 100949. 10.1016/J.MCPRO.2025.100949.

24. Bubis, J.A., Arrey, T.N., Damoc, E., Delanghe, B., Slovakova, J., Sommer, T.M., Kagawa, H., Pichler, P., Rivron, N., Mechtler, K., et al. (2025). Challenging the Astral mass analyzer to quantify up to 5,300 proteins per single cell at unseen accuracy to uncover cellular heterogeneity. Nature Methods 2025 22:3 **22**, 510–519. 10.1038/s41592-024-02559-1.

25. Sydney Shaffer, A.M., Emert, B.L., Reyes Hueros, R.A., Singh, A., Bassett, D.S., and Raj Correspondence, A. (2020). Memory Sequencing Reveals Heritable Single-Cell Gene Expression Programs Associated with Distinct Cellular Behaviors. Cell 182, 947–959.e17. 10.1016/j.cell.2020.07.003.

26. Mold, J.E., Weissman, M.H., Ratz, M., Hagemann-Jensen, M., Hård, J., Eriksson, C.-J., Toosi, H., Berghenstråhle, J., Berlin, L. von, Martin, M., et al. (2022). Clonally heritable gene expression imparts a layer of diversity within cell types. bioRxiv, 2022.02.14.480352. 10.1016/j.cels.2024.01.004.

27. Sigal, A., Milo, R., Cohen, A., Geva-Zatorsky, N., Klein, Y., Liron, Y., Rosenfeld, N., Danon, T., Perzov, N., and Alon, U. (2006). Variability and memory of protein levels in human cells. Nature 444, 643–646. 10.1038/nature05316.

28. Spencer, S.L., Gaudet, S., Albeck, J.G., Burke, J.M., and Sorger, P.K. (2009). Non-genetic origins of cell-to-cell variability in TRAIL-induced apoptosis. Nature 459, 428–432. 10.1038/nature08012.

29. PUCK, T.T., and MARCUS, P.I. (1956). Action of x-rays on mammalian cells. Volume 103, Issue 5, Pages 653 - 666 **103**, 653–666. 10.1084/jem.103.5.653.

30. Brix, N., Samaga, D., Belka, C., Zitzelsberger, H., and Lauber, K. (2021). Analysis of clonogenic growth in vitro. Nat. Protoc. 16, 4963–4991. 10.1038/S41596-021-00615-0.

31. Rangan, S.R.S. (1972). A new human cell line (FaDu) from a hypopharyngeal carcinoma. Cancer 29, 117–121. 10.1002/1097-0142(197201)29:1<117::AID-CNCR2820290119>3.0.CO;2-R.

32. Piga, I ., Koenig, C., Lechner, M., Sabatier, P., and Olsen, J. V. (2025). Formaldehyde Fixation Helps Preserve the Proteome State during Single-Cell Proteomics Sample Processing and Analysis. J. Proteome Res. 24, 1624–1635. 10.1021/ACS.JPROTEOME.4C00656/ASSET/IMAGES/LARGE/PR4C00656_0006.JPEG.

33. Meier, F., Brunner, A.D., Frank, M., Ha, A., Bludau, I ., Voytik, E., Kaspar-Schoenefeld, S., Lubeck, M., Raether, O., Bache, N., et al. (2020). diaPASEF: parallel accumulation–serial fragmentation combined with data-independent acquisition. Nat. Methods 17, 1229–1236. 10.1038/s41592-020-00998-0.

34. Demichev, V., Messner, C.B., Vernardis, S.I ., Lilley, K.S., and Ralser, M. (2020). DIA-NN: neural networks and interference correction enable deep proteome coverage in high throughput. Nat. Methods 17, 41–44. 10.1038/s41592-019-0638-x.

35. Ly, T., Whigham, A., Clarke, R., Brenes-Murillo, A.J., Estes, B., Madhessian, D., Lundberg, E., Wadsworth, P., and Lamond, A.I. (2017). Proteomic analysis of cell cycle progression in asynchronous cultures, including mitotic subphases, using PRIMMUS. Elife 6, 1–35. 10.7554/eLife.27574.

36. Bubis, J.A., Arrey, T.N., Damoc, E., Delanghe, B., Slovakova, J., Sommer, T.M., Kagawa, H., Pichler, P., Rivron, N.C., Mechtler, K., et al. (2024). Challenging the Astral mass analyzer - up to 5300 proteins per single-cell at unseen quantitative accuracy to study cellular heterogeneity. bioRxiv, 2024.02.01.578358. 10.1038/s41592-024-02559-1.

37. Matsunuma, R., Chan, D.W., Kim, B.J., Singh, P., Han, A., Saltzman, A.B., Cheng, C., Lei, J.T., Wang, J., Roberto da Silva, L., et al. (2018). DPYSL3 modulates mitosis, migration, and epithelial-to-mesenchymal transition in claudin-low breast cancer. Proc. Natl. Acad. Sci. U. S. A. 115, E11978–E11987. 10.1073/PNAS.1810598115/SUPPL_FILE/PNAS.1810598115.SD03.XLSX.

38. Tsherniak, A., Vazquez, F., Montgomery, P.G., Weir, B.A., Kryukov, G., Cowley, G.S., Gill, S., Harrington, W.F., Pantel, S., Krill-Burger, J.M., et al. (2017). Defining a Cancer Dependency Map. Cell 170, 564–576.e16. 10.1016/J.CELL.2017.06.010.

39. Johnson, D.E., Burtness, B., Leemans, C.R., Lui, V.W.Y., Bauman, J.E., and Grandis, J.R. (2020). Head and neck squamous cell carcinoma. Nat. Rev. Dis. Primers 6. 10.1038/S41572-020-00224-3.

40. Nör, C., Zhang, Z., Warner, K.A., Bernardi, L., Visioli, F., Helman, J.I ., Roesler, R., and Nör, J.E. (2014). Cisplatin induces Bmi-1 and enhances the stem cell fraction in head and neck cancer. Neoplasia (United States) 16, 137–146. 10.1593/neo.131744.

41. Sharma, A., Cao, E.Y., Kumar, V., Zhang, X., Leong, H.S., Wong, A.M.L., Ramakrishnan, N., Hakimullah, M., Teo, H.M.V., Chong, F.T., et al. (2018). Longitudinal single-cell RNA sequencing of patient-derived primary cells reveals drug-induced infidelity in stem cell hierarchy. Nat. Commun. 9. 10.1038/S41467-018-07261-3.

42. Niehr, F., Eder, T., Pilz, T., Konschak, R., Treue, D., Klauschen, F., Bockmayr, M., Türkmen, S., Jöhrens, K., Budach, V., et al. (2018). Multilayered omics-based analysis of a head and neck cancer model of cisplatin resistance reveals intratumoral heterogeneity and treatment-induced clonal selection. Clinical Cancer Research 24, 158–168. 10.1158/1078-0432.CCR-17-2410.

43. Niehr, F., Eder, T., Pilz, T., Konschak, R., Treue, D., Klauschen, F., Bockmayr, M., Türkmen, S., Jöhrens, K., Budach, V., et al. (2018). Multilayered omics-based analysis of a head and neck cancer model of cisplatin resistance reveals intratumoral heterogeneity and treatment-induced clonal selection. Clinical Cancer Research 24, 158–168. 10.1158/1078-0432.CCR-17-2410.

44. Tinhofer, I ., Budach, V., Saki, M., Konschak, R., Niehr, F., Jöhrens, K., Weichert, W., Linge, A., Lohaus, F., Krause, M., et al. (2016). Targeted next-generation sequencing of locally advanced squamous cell carcinomas of the head and neck reveals druggable targets for improving adjuvant chemoradiation. Eur. J. Cancer 57, 78–86. 10.1016/j.ejca.2016.01.003.

45. Tanaka, N., Zhao, • Mei, Lin Tang, •, Patel, A.A., Xi, Q., Van, H.T., Takahashi, H., Osman, A.A., Zhang, J., Wang, J., et al. (2018). Gain-of-function mutant p53 promotes the oncogenic potential of head and neck squamous cell carcinoma cells by targeting the transcription factors FOXO3a and FOXM1. Oncogene 37, 1279–1292. 10.1038/s41388-017-0032-z.

46. Amlani, L., Bellile, E., Spector, M., Smith, J., Brenner, C., Rozek, L., Nguyen, A., Zarins, K., Thomas, D., McHugh, J., et al. (2019). Expression of P53 and Prognosis in Patients with Head and Neck Squamous Cell Carcinoma (HNSCC). Int. J. Cancer Clin. Res. 6. 10.23937/2378-3419/1410122.

47. Boysen, M., Kityk, R., and Mayer, M.P. (2019). Hsp70- and Hsp90-Mediated Regulation of the Conformation of p53 DNA Binding Domain and p53 Cancer Variants. Mol. Cell 74, 831–843.e4. 10.1016/J.MOLCEL.2019.03.032/ATTACHMENT/23CCCDC6-BEA6-4D92-80A7-0ADA4D7396A0/MMC2.PDF.

48. Zylicz, M., King, F.W., and Wawrzynow, A. (2001). Hsp70 interactions with the p53 tumour suppressor protein. EMBO J. 20, 4634–4638. 10.1093/EMBOJ/20.17.4634.

49. Barkley, D., Moncada, R., Pour, M., Liberman, D.A., Dryg, I ., Werba, G., Wang, W., Baron, M., Rao, A., Xia, B., et al. (2022). Cancer cell states recur across tumor types and form specific interactions with the tumor microenvironment. Nature Genetics 2022 54:8 **54**, 1192–1201. 10.1038/s41588-022-01141-9.

50. Weichselbaum, R.R., Ishwaran, H., Yoon, T., Nuyten, D.S.A., Baker, S.W., Khodarev, N., Su, A.W., Shaikh, A.Y., Roach, P., Kreike, B., et al. (2008). An interferon-related gene signature for DNA damage resistance is a predictive marker for chemotherapy and radiation for breast cancer. Proc. Natl. Acad. Sci. U. S. A. 105, 18490–18495. 10.1073/pnas.0809242105.

51. Salovska, B., Li, W., Bernhardt, O.M., Germain, P.L., Wang, Q., Gandhi, T., Reiter, L., and Liu, Y. (2025). A robust multiplex-DIA workflow profiles protein turnover regulations associated with cisplatin resistance and aneuploidy. Nature Communications 2025 16:1 **16**, 5034-. 10.1038/s41467-025-60319-x.

52. Weber, K., Thomaschewski, M., Warlich, M., Volz, T., Cornils, K., Niebuhr, B., Täger, M., Lütgehetmann, M., Pollok, J.M., Stocking, C., et al. (2011). RGB marking facilitates multicolor clonal cell tracking. Nature Medicine 2011 17:4 **17**, 504–509. 10.1038/nm.2338.

53. Cornils, K., Thielecke, L., Hüser, S., Forgber, M., Thomaschewski, M., Kleist, N., Hussein, K., Riecken, K., Volz, T., Gerdes, S., et al. (2014). Multiplexing clonality: Combining RGB marking and genetic barcoding. Nucleic Acids Res. 42. 10.1093/nar/gku081.

54. Wood, R.D. (1997). Nucleotide excision repair in mammalian cells. Journal of Biological Chemistry 272, 23465–23468. 10.1074/jbc.272.38.23465.

55. Lemaire, M.A., Schwartz, A., Rahmouni, A.R., and Leng, M. (1991). Interstrand cross-links are preferentially formed at the d(GC) sites in the reaction between cis-diamminedichloroplatinum (II) and DNA. Proceedings of the National Academy of Sciences 88, 1982–1985. 10.1073/PNAS.88.5.1982.

56. Zhou, H., Zhao, C., Shao, R., Xu, Y., and Zhao, W. (2023). The functions and regulatory pathways of S100A8/A9 and its receptors in cancers. Front. Pharmacol. 14, 1187741. 10.3389/FPHAR.2023.1187741.

57. Argyris, P.P., Slama, Z., Malz, C., Koutlas, I .G., Pakzad, B., Patel, K., Kademani, D., Khammanivong, A., and Herzberg, M.C. (2019). Intracellular calprotectin (S100A8/A9) controls epithelial differentiation and caspase-mediated cleavage of EGFR in head and neck squamous cell carcinoma. Oral Oncol. 95, 1–10. 10.1016/J.ORALONCOLOGY.2019.05.027.

58. Mund, A., Coscia, F., Kriston, A., Hollandi, R., Kovács, F., Brunner, A.D., Migh, E., Schweizer, L., Santos, A., Bzorek, M., et al. (2022). Deep Visual Proteomics defines single-cell identity and heterogeneity. Nature Biotechnology 2022 40:8 40, 1231–1240. 10.1038/s41587-022-01302-5.

59. Vizcaíno, J.A., Csordas, A., Del-Toro, N., Dianes, J.A., Griss, J., Lavidas, I., Mayer, G., Perez-Riverol, Y., Reisinger, F., Ternent, T., et al. (2016). 2016 update of the PRIDE database and its related tools. Nucleic Acids Res. 44, D447–56. 10.1093/nar/gkv1145.

60. Fisch, A.S., Pestana, A., Sachse, V., Doll, C., Hofmann, E., Heiland, M., Obermueller, T., Heidemann, J., Dommerich, S., Schoppe, D., et al. (2024). Feasibility analysis of using patient-derived tumour organoids for treatment decision guidance in locally advanced head and neck squamous cell carcinoma. Eur. J. Cancer 213. 10.1016/j.ejca.2024.115100.

61. Makhmut, A., Qin, D., Fritzsche, S., Nimo, J., König, J., and Coscia, F. (2023). A framework for ultra-low-input spatial tissue proteomics. Cell Syst. 14, 1002–1014.e5. 10.1016/j.cels.2023.10.003.

62. Ctortecka, C., Clark, N.M., Boyle, B.W., Seth, A., Mani, D.R., Udeshi, N.D., and Carr, S.A. (2024). Automated single-cell proteomics providing sufficient proteome depth to study complex biology beyond cell type classifications. Nat. Commun. 15, 1–10. 10.1038/S41467-024-49651-W;TECHMETA.

